# Light sensitive short hypocotyl (LSH) confer symbiotic nodule identity in the legume Medicago truncatula

**DOI:** 10.1101/2023.02.12.528179

**Authors:** K. Schiessl, T. Lee, M. Orvosova, M. Bueno-Batista, N. Stuer, P.C. Bailey, K.S. Mysore, J. Wen, G.E.D Oldroyd

## Abstract

Legumes grow specialized root nodules that are distinct from lateral roots in morphology and function, with nodules intracellularly hosting beneficial nitrogen-fixing bacteria that provide the plant with a nitrogen source. We have previously shown that a lateral root-like program underpins nodule initiation, but there must be additional developmental programs that confer nodule identity. Here, we show that two members of the *LIGHT SENSITIVE SHORT HYPOCOTYL (LSH)* transcription factor family, predominantly known to define organ boundaries and meristem complexity in the shoot, function as regulators of nodule organ identity. *LSH1/LSH2* function upstream of and together with the known nodule regulators *Nuclear Factor Y*-*A1* and *NODULE ROOT1/2.* The principal outcome of *LSH1/LSH2* function is the production of cells able to accommodate nitrogen-fixing bacteria, the unique nodule feature. We conclude that the coordinate recruitment of a pre-existing primordium identity program, in parallel to a root initiation program, underpins the divergence between lateral roots and nodules.

## Introduction

To overcome nitrogen deficiencies in the soil, legumes have evolved the ability to enter symbioses with beneficial nitrogen-fixing rhizobial bacteria. Legumes accommodate rhizobia inside cells of specialized root organs called nodules, that provide a favorable environment for N-fixation. Establishing this interaction involves legume recognition of rhizobial Nod factors by plant receptor-like kinases at the root surface, that promote symbiosis signaling (Oldroyd, 2013). This in turn initiates two spatially separated but temporally coordinated processes. At root epidermal cells, the attached rhizobial bacteria are entrapped in a root hair curl and invade the cell via infection threads, tip-growing plasma membrane invaginations. Concomitantly, the development of the nodule organ primordium is initiated via cell divisions in the inner tissue layers of the root below the site of infection. As the infection thread progresses and meets the nodule primordium, the bacteria are released and internalized into membrane-bound compartments, the symbiosomes, where they transition into the nitrogen-fixing state (Oldroyd et al., 2011; Roy et al., 2020).

Initiation and progression of bacterial infection and nodule development are controlled by and dependent on the function of the nodulation-specific transcriptional regulator *NODULE INCEPTION* (*NIN*), which is activated downstream of the symbiosis signaling pathway (Marsh et al., 2007; Singh et al., 2014). Epidermal infection and nodule organogenesis are genetically separable processes as evidenced by studies of different *nin* loss of function alleles and studies of the *cis* regulatory elements in the *NIN* promoter (Liu et al., 2021; Liu et al., 2019; Marsh et al., 2007; Schauser et al., 1999; Vernie et al., 2015; Yoro et al., 2014). To initiate the development of symbiotic root nodules in the inner tissue layers, *NIN* expression is activated via *CYTOKININ RESPONSE 1* (*CRE1*)-mediated cytokinin signaling (Gonzalez-Rizzo et al., 2006; Liu et al., 2019). Our previous work showed that cytokinin-induced *NIN* recruits a conserved program associated with lateral root development, to initiate the formation of a symbiotic root nodule, through the transcriptional regulator *LATERAL ORGAN BOUNDARIES 16* (*LBD16*), which promotes cell proliferation in the inner tissue layers via the upregulation of *STYLISH-like* (*STY-l*) transcriptional regulators and *YUCCA* (*YUC*) auxin biosynthesis genes (Schiessl et al., 2019). Consequently, nodules and lateral roots initiate from the same inner tissue layers of the primary root including the pericycle, the endodermis and the inner cortex in response to the local accumulation of auxin (Herrbach et al., 2014; Schiessl et al., 2019).

The commonalities between lateral root and nodule initiation imply that additional nodule-specific regulatory pathways must be recruited to confer nodule organ identity, in order to differentiate nodules from lateral roots in both morphology and function. While lateral root primordia predominantly develop from cells that are derived from the inner cell layers, the development of nodule primordia is specifically associated with the promotion of cell proliferation in the mid-cortex (Herrbach et al., 2014; Xiao et al., 2014). These cortical-derived primordium cells are key determinants in the establishment of nodule organ identity as they undergo host cell differentiation and are intracellularly colonized by the rhizobial bacteria that are released from the infection thread (Xiao et al., 2014).

Several regulators have been implicated in the promotion of cortical cell divisions downstream of cytokinin-induced *NIN* expression. These include the transcriptional regulator *NUCLEAR FACTOR Y* (*NF-YA1*) and the *SHORT ROOT–SCARECROW* (*SHR-SCR*) stem cell regulatory module (Dong et al., 2021; Laloum et al., 2014; Xiao et al., 2014). *NF-YA1* functions in cortical infection thread progression, nodule initiation, and the subsequent establishment and maintenance of the nodule meristem, which provides cells for continued rhizobial infection in the indeterminate nodule of *M. truncatula* (Combier et al., 2006; Soyano et al., 2013; Soyano et al., 2019; Xiao et al., 2014). *NF-YA1* does so in part through the upregulation of *STY* transcription factors and *YUC* auxin biosynthesis genes (Hossain et al., 2016; Schiessl et al., 2019; Shrestha et al., 2020). *LBD16* induction by *NIN* initiates a root-like program, which appears to be later suppressed by *NODULE ROOT1/2* (*NOOT1/NOOT2*), mutations of which develop root-like conversions from nodules (Magne et al., 2018). *NOOT1/NOOT2* are orthologous to the *Arabidopsis* genes *BLADE-ON-PETIOLE 1/2* (*BOP1/2*), two BTB-ankyrin transcriptional co-activators with multiple functions in the regulation of lateral organ development in the shoot (Couzigou et al., 2012; Khan et al., 2014; Magne et al., 2018). While nodule development is compromised in *nf-ya1* and *noot1/noot2*, nodules still initiate in these mutants, that can at least in part support rhizobial colonization and N-fixation, suggesting that additional regulators of nodule organ identity must act either upstream, redundantly or in parallel with these known regulators. In this study, we demonstrate two members of the *LIGHT SENSITIVE SHORT HYPOCOTYL (LSH)* transcription factor family, that previously have been shown to predominantly control shoot meristem function, also control nodule organ identity, downstream of *NIN.* These genes are required to provide the unique organ identity of nodules, allowing intracellular bacterial colonization and N-fixation, through the upregulation of *NF-YA1* and *NOOT1/NOOT2,* together with additional shoot-expressed developmental regulators. The coordinated recruitment of pre-existing growth regulators and a root developmental program appears to allow initiation of root nodules and their unique specification.

## RESULTS

### *LSH1* and *LSH2* are upregulated during early nodule organogenesis downstream of *NIN*

To define potential regulators of nodule organ identity we screened a pre-existing comparative RNA-Seq time course of nodule and lateral root development, for those induced during the early stages of nodule development, with nodule-specific expression, in a manner dependent on *NIN* (Feng et al., 2021; Schiessl et al., 2019). This selection criteria identified *NF-YA1*, its interacting subunit *NF-YB16* and *NOOT1*/*NOOT2* (Baudin et al., 2015; Magne et al., 2018; Soyano et al., 2013), as well as two transcriptional regulators with yet uncharacterized functions in the rhizobial symbiosis (Figure 1A). These novel regulators share an ALOG domain of identical amino acid sequence, with high sequence similarity to *LIGHT SENSITIVE SHORT HYPOCOTYL* (*LSH*) transcription factor 3 in *Arabidopsis* (*AtLSH3*) and with the recently characterized *SYMMETRIC PETALS 1* gene in *Pisum sativum* (*PsSYP1*) (Figure S1A) (He et al., 2020; Iyer and Aravind, 2012; Naramoto et al., 2020). We named these two genes *LSH1* and *LSH2*. *LSH1* is upregulated in roots from 16 hrs post rhizobial spot inoculation, while *LSH2* is upregulated from 36 hrs. By contrast, neither *LSH1* or *LSH2* were differentially expressed during lateral root development, suggesting that *LSH1* and *LSH2* may be part of a developmental program that distinguishes nodules from lateral roots (Figure 1A) (Schiessl et al., 2019). The expression of *LSH1* and *LSH2* during rhizobial infection was dependent on *CRE1* and *NIN*, and ectopic expression of *NIN* was sufficient to upregulate both genes (Figure 1A) (Feng et al., 2021; Schiessl et al., 2019). Furthermore, expression of *LSH1* was induced by cytokinin treatment of *M. truncatula* roots in a *CRE1-* and *NIN-*dependent but *NF-YA1*-independent manner (Figure 1B). To investigate their spatial expression patterns, we performed promoter-GUS analysis of *LSH1* and *LSH2,* which revealed that both genes are expressed in newly divided cells of the nodule primordia throughout development. In mature nodules, *LSH1* is expressed in the apical meristem region and *LSH2* is expressed in the infection and fixation zones, with both genes expressed in the peripheral nodule vasculature (Figures 1C and S1B).

**Figure 1:**
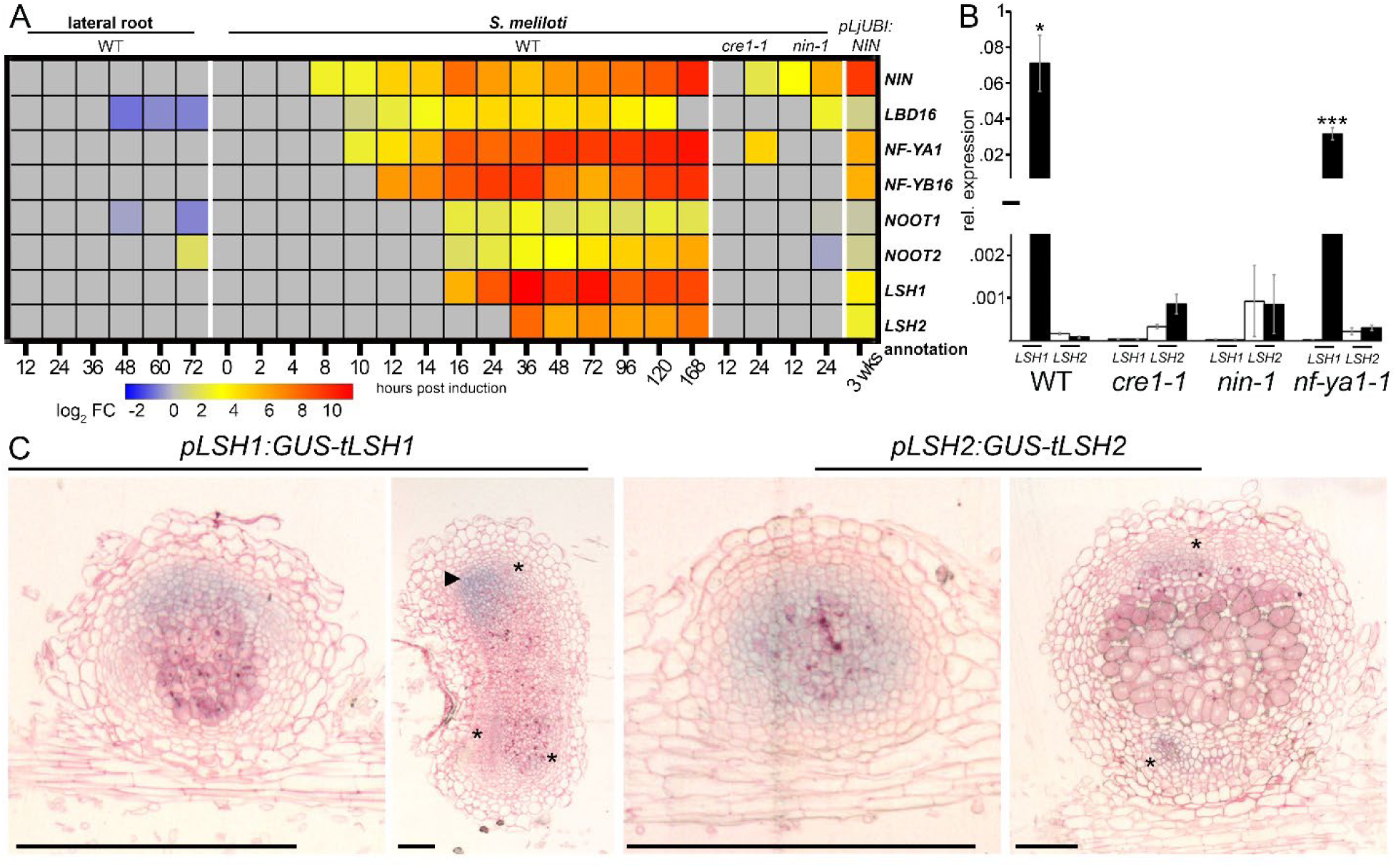
*LSH1* and *LSH2* are upregulated during early nodule organogenesis downstream of *NIN*. (A) Heatmap shows selected genes induced during lateral root and nodule development. Fold changes compared to controls are depicted in log2 scale with the significance threshold of p-value < 0.05. (B) Expression profiling on root segments treated with 100 nM 6-Benzylaminopurine (BAP) for 24 h by qRT-PCR normalized to *HH3*. Statistical comparisons between mock (white bars) and BAP (black bars). Values are the mean ΔCt values of 3 biological replicates for *LSH1* and 2 for *LSH2* ± SEM (Student’s t-test; asterisks indicate statistical significance *, P < 0.05; **, P < 0.01, ***, P < 0.001). (C) Expression patterns of *LSH1* and *LSH2* visualized by GUS staining (blue) (also see Figure S1B). Rhizobial expressed *LacZ* is stained magenta. Ruthenium Red demarks cell walls. Asterisks indicate expression in vascular bundles and arrows in the meristem. Scale bars: 500 µm.

### *LSH1/LSH2* are required for nodule development and N- fixation

To assess the role of *LSH1* and *LSH2* during nodule organogenesis, we identified loss of function mutants in *LSH1 (lsh1-1, lsh1-2)* and *LSH2 (lsh2-1)* and generated an *lsh1-1/lsh2-1* double mutant (Figure S2A). *lsh1,* but not *lsh2*, showed significantly altered shoot organ morphologies including changes in petal shape and number, and a reduction in stipule complexity (Figures 2A-B and S2B and D, Table S1), showing conserved functions for this gene across many angiosperms (Cho and Zambryski, 2011; He et al., 2020; MacAlister et al., 2012; Yoshida et al., 2013). No effects of *LSH* have been reported on root system architecture, and we also found that the root morphology was unaffected in lsh1/lsh2 seedlings and that *LSH1/LSH2* are not positive regulators of lateral root development. In fact we observed a slight increase in the number of lateral roots in *lsh1* and *lsh1/lsh2* seedlings besides a slight reduction in primary root length (Figure S2C), suggesting that in *M. truncatula* these genes are negative regulators of lateral root initiation.

**Figure 2:**
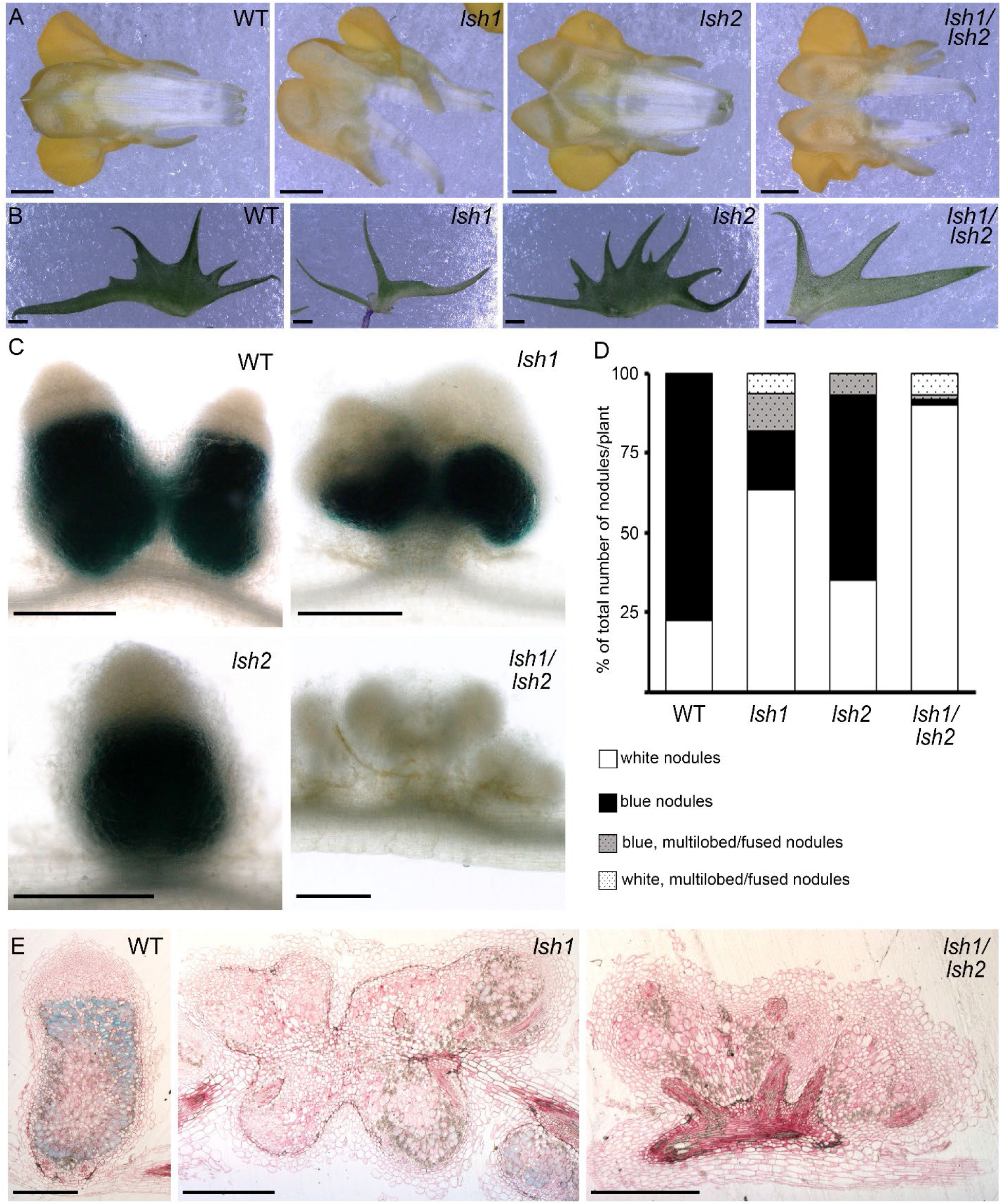
*LSH1/LSH2* are required for nodule development and N-fixation. (A-B) Images of WT, *lsh1-2*, *lsh2-1* and *lsh1-1/lsh2-1* dissected flower keels (A) and stipules (B), also see Figures S2B and D, Table S1). Scale bars: 500 µm. (C) Whole mount images of WT, *lsh1-1*, *lsh2-1* and *lsh1-1/lsh2-1* nodules 28 days post *S. meliloti* inoculation. GUS staining (blue) indicates the expression of the bacterial *pNifH* promoter. Scale bars: 500 µm. (D) Distribution of different nodule morphologies depicted as percentage of total nodule number per plant in WT and *lsh1-1* (n = 15), *lsh2-1* (n = 14), and *lsh1-1/lsh2-1* (n = 13) (also see Figure S2E-F). (E) Sections of 28-d old nodules in WT, *lsh1-1* and *lsh1-1/lsh2-1, pNifH:GUS* expression (blue), Ruthenium Red demarks cell walls. Scale bars: 500 µm.

For assessment of functions during nodulation, we inoculated plants with *Sinorhizobium meliloti* strain 2011 expressing *pNifH-GUS*, a bacterial promoter associated with the expression and activity of nitrogenase and used to approximate biological N-fixation in nodules (Starker et al., 2006). *lsh1* mutants showed a high frequency of white nodules, while the *lsh1/lsh2* double mutant developed almost exclusively white nodules, suggesting that N-fixation was severely attenuated (Figures 2C-D and S2E-F). The reduction and absence of N-fixation was confirmed in an acetylene reduction assay (Figure S2G). *lsh1/lsh2* showed an increase in nodule number (Figure S2H), a phenomenon frequently observed in fix^-^ mutants. Nodule morphology was significantly altered in the *lsh1* and *lsh1/lsh2*, with multilobed and fused nodules (Figures 2C-E and S2E-F). Nodule sections revealed a severe reduction in the number of cells with intracellular bacterial colonization in *lsh1/lsh2* nodules, but no obvious defects were observed in bacteroid differentiation in the few cells that possessed intracellular colonization (Figure S3A).

### *LSH* genes are required for the development of nodule primordia that can support bacterial colonization

To further investigate the cause of this severe reduction in rhizobial colonisation, we assessed the progression of rhizobial infection and early nodule primordium development. In wild type, the progression of rhizobial infection threads through the epidermis and cortex was temporally and spatially coordinated with the development of the nodule primordium (Figures 3A-B and S6A-B). We observed nodule primordium initiation and early establishment of epidermal infection threads in *lsh1* and *lsh1/lsh2*, however, the progression of infection threads through the cortex and subsequent internal colonization of primordium cells was severely impaired (Figures 3A-B and S6A-B). This resulted in a high proportion of *lsh1* and *lsh1/lsh2* nodules that were only partially or completely uncolonized, at the point of emergence from the primary root.

**Figure 3.**
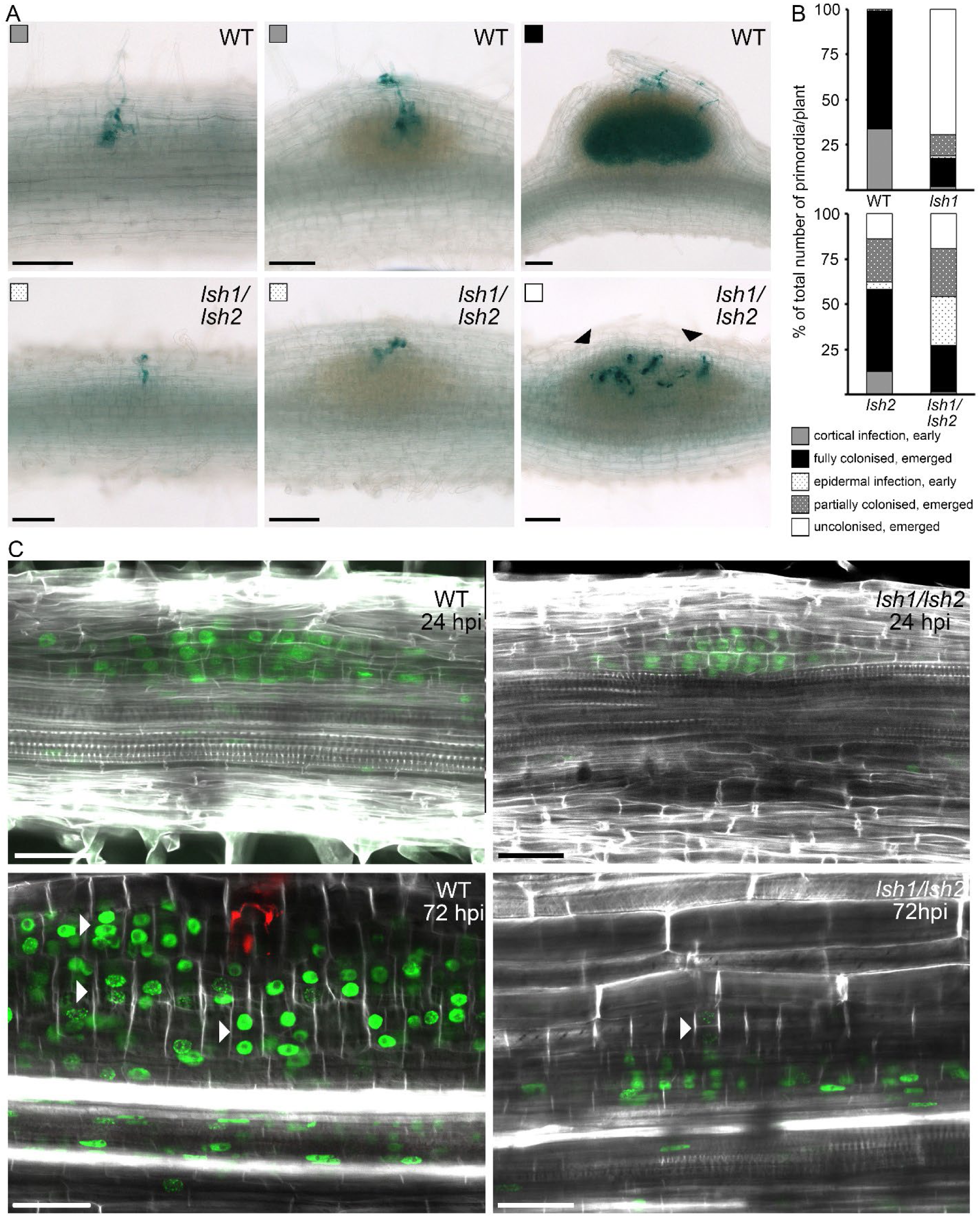
*LSH* genes are required for the development of nodule primordia that can support bacterial colonization. (A) Images of WT and *lsh1/lsh2* nodule primordia at different developmental stages, first initial divisions (left), multilayered (middle) and emerged primordia (right) observed 7 d post spray inoculation with rhizobial bacteria expressing *LacZ* (blue stain). Black arrowheads indicate infection threads that are restricted in their progression into the inner root tissue layers. Squares relate to the legend in Figure 3B. Scale bars: 500 µm. (B) Distribution of bacterial colonization phenotypes observed in WT, *lsh1*, *lsh2* and *lsh1/lsh2* at 7 dpi depicted as percentage of the total primordia number per plant, WT (n = 30), *lsh1, lsh2* (n = 33), *lsh1/lsh2* (n = 32). (C) Optical sections of WT and *lsh1/lsh2* roots 24 and 72 h post spot-inoculation with *S. meliloti* (n >30 per genotype and timepoint), also see Figure S3B. At 72 hpi, *Sm2011-mCherry* bacteria in red, cell walls in white (fluorescent brightener) and EdU-labelled nuclei indicating DNA replication in green. White arrowheads indicate periclinal cell divisions. Scale bars: 50 µm.

To study the early development of the nodule primordia, we used rhizobial spot inoculation combined with deep tissue imaging using the DNA synthesis marker 5-ethynyl-2-deoxyuridine (EdU) combined with either propidium iodide or fluorescent brightener as a cell wall marker (Schiessl et al., 2019; Ursache et al., 2018). At 24 hrs post rhizobial spot inoculation (24 hpi), we observed no differences in the cell cycle activity and overall primordium development between WT and *lsh1/lsh2*, but at 72 hpi we observed a severe reduction in cell cycle activity in *lsh1/lsh2* primordia (Figures 3C and S3B). This reduction in cell cycle activity appeared to be specific to the primordium cells derived from the middle cortex of the primary root, while the cell cycle activity in the inner tissue layers at the base of the primordia appeared comparable between WT and *lsh1/lsh2*. WT primordia underwent periclinal cell divisions at 72 hpi, adding cortical derived cell layers to the growing primordium, which were absent in *lsh1/lsh2* (Figures 3C and S3B). We conclude that *LSH1*/*LSH2* are not required for the early initiation of the nodule primordium, but rather play a role in divisions of the cortical cell layer, that give rise to cells that accommodate the N-fixing bacteria (Xiao et al., 2014).

### *LSH1* and *LSH2* are required for the upregulation of nodule organ identity genes and the recruitment of known growth regulators during nodule organogenesis

In order to better understand the regulatory function of the transcription factors *LSH1* and *LSH2* during rhizobial infection and nodule organogenesis, we performed RNA-Seq on rhizobial spot inoculated root sections of *lsh1* and *lsh1/lsh2* at 24 and 72 hpi. In addition, we generated hairy roots expressing both *pLjUBI:LSH1* and *pLjUBI:LSH2* and performed RNA-Seq under non-symbiotic conditions. Consistent with an early role in nodule primordium formation, *lsh1* and *lsh1/lsh2* showed severe reductions in nodule-associated gene expression compared to wild type, with over 90% of rhizobial-responsive genes not differentially expressed in the *lsh1/2* mutant. (Figure 4A, Data S1). Marker genes for symbiosis signaling, such as *EARLY NODULIN 11 (ENOD11)* and *NIN*, were still expressed in *lsh1/lsh2*, as were genes associated with early infection such as *Nodule Pectate Lyase* (*NPL*) and *Rhizobium-directed Polar Growth (RPG)* (Figure 4B). In contrast, genes associated with infection progression and N-fixation such as *VAPYRIN (VYP)* and *LEGHEMOGLOBIN*s (*LB1*/*LB2*) were not upregulated in the *lsh1/lsh2* mutant (Figure 4B). Of the genes known to be associated with the initiation of the nodule primordium, *LBD16* expression was not significantly affected by *LSH* mutation, whereas the known nodule regulators *NF-YA1* and *NOOT1/NOOT2* showed partial dependency on *LSH1/LSH2* in the loss and gain of function context. *NF-YA1* has been shown to regulate the expression of *STY-l* transcription factors, which in turn promote expression of the *YUC* auxin biosynthesis genes (Hossain et al., 2016; Shrestha et al., 2020) and consistently *STY-l* and *YUC* genes were downregulated in the *lsh* mutants. Related to this, we found that several genes involved in auxin transport and conjugation, but also cell cycle regulators including A-type and B-type cyclins and the endoreduplication regulator *CSS52B* were also dependent on *LSH1*/*LSH2.* Many of these genes were constitutively upregulated by overexpression of *LSH1/LSH2* (Figure 4B).

**Figure 4.**
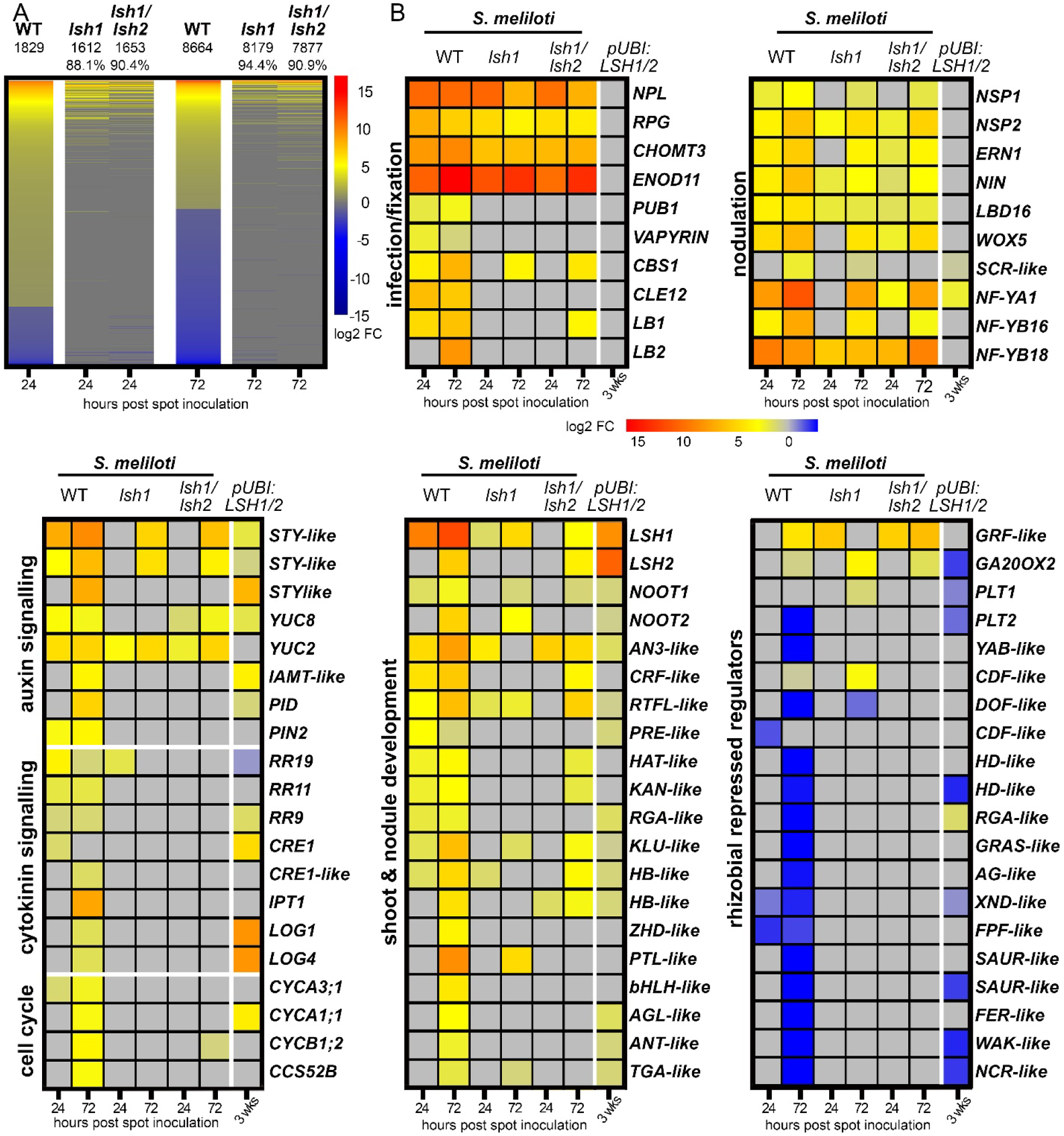
*LSH1* and *LSH2* are required for the upregulation of nodule organ identity genes and the recruitment of shoot-expressed genes during nodule organogenesis. (A) Heatmaps of all differentially expressed genes (DEGs) in response to *S. meliloti* spot inoculation in WT and *lsh1* and *lsh1/lsh2* at 24 and 72 hpi. Expression levels are depicted as log_2_ fold changes (log_2_ fold changes ≥ +/-1, p-value < 0.05). To compare the overall transcriptional response to *S. meliloti* spot inoculation between WT and the mutants, we sorted all DEGs from the highest positive to the highest negative log_2_ fold change value. Absolute numbers indicate the number of DEGs in each genotype. Percentages indicate the proportion of DEGs in WT that are not expressed in the mutants and therefore dependent on *LSH1* and *LSH1/2*. (B) Heatmaps showing expression levels of selected functional groups of DEGs in WT, *lsh1* and *lsh1/lsh2* root sections at 24 and 72 hpi and in response to combined ectopic expression of *LSH1*/*LSH2* (*pUBI:LSH1/LSH2*) compared to empty vector in 3-week old WT (jemalong) hairy roots. Fold changes compared to controls are depicted in log2 scale with the significance threshold of p-value < 0.05. Related to Figures 5C, 6B-C and S5A-C. Also see Data S1.

Cytokinin signaling is necessary and sufficient for nodule initiation and development (Cooper and Long, 1994; Gauthier-Coles et al., 2019; Gonzalez-Rizzo et al., 2006; Murray et al., 2007; Tirichine et al., 2007)) and has also been shown to be required for endosymbiotic host cell colonization by facilitating the switch from mitotic cell proliferation to endoreduplication through the upregulation of *CSS52A* (Tan et al., 2020). *LSH1* and *LSH2* induction by rhizobia requires *CRE1* and *NIN* (Figure 1A), suggesting they act downstream of cytokinin promotion of *NIN* in the root cortex. Surprisingly, however, we found that genes with a function in cytokinin signaling and biosynthesis were strongly affected by the loss and gain of *LSH1/LSH2* function, including *CRE1*, the *B-type RESPONSE REGULATORS RR9* and *RR11* and members of the *LONELY GUY (LOG)* gene family (Kuroha et al., 2009; Mortier et al., 2014) (Figure 4B). This promotion of cytokinin signaling and its mode of action may provide positive feedback to further drive nodule organ identity.

Previously, members of the *LSH* transcription factor family have been characterized to function together with the transcriptional co-activators *BLADE-ON-PETIOLE* in the shoot where they control the complexity of inflorescences and asymmetry of floral organs (He et al., 2020; Xu et al., 2016). Consistently, we found a subset of developmental regulators that were upregulated in response to rhizobial spot inoculation in an *LSH*-dependent manner and also found to be expressed in shoot tissues of *M. truncatula* (data extracted from the MtExpress *Medicago* Gene expression atlas; (Carrere et al., 2021)). Several of these *LSH*-dependent and shoot-expressed genes have been previously annotated to function as regulators of organ growth and organ boundaries such as *KLUH* and *PETAL LOSS* (Anastasiou et al., 2007; Brewer et al., 2004). We also found a set of rhizobial-repressed genes that were *LSH1/LSH2*-dependent, including two members of the *PLETHORA (PLT)* root meristem regulator family, *PLT1* and *PLT2* (Franssen et al., 2015). Together, our RNA-Seq analyses demonstrate the *LSH* genes as major regulators of nodulation that are necessary and sufficient for the up-regulation of nodule-specific regulators such as *NF-YA1* and the recruitment of regulators with pleiotropic functions in shoot and symbiotic nodule development, including *NOOT1/NOOT2*.

### *LSH1/LSH2* partly functions through the cortical activation of *NF-YA1*

To understand the spatial context of *LSH1/LSH2* function, we performed promoter GUS analysis in hairy roots expressing *pNF-YA1:GUS-tNF-YA1* in wild-type and *lsh1/lsh2* background. *pNF- YA1:GUS-tNF-YA1* showed expression in wild type in the inner tissue layers at the base of the developing nodule and in the nodule primordium (Figures 5A). In the *lsh1/lsh2* mutant we observed loss of *NF-YA1* expression in the nodule primordium, but the maintenance of its expression in the tissue layers at the base of the nodule (Figures 5A). Such tissue-specific control of *NF-YA1* is consistent with the partial *LSH1/LSH2* dependency for *NF-YA1* induction observed in the RNA-Seq (Figure 4B). Previously, *NF-YA1* has been characterized to play a crucial role in promoting cell proliferation, host cell differentiation and endosymbiotic colonization in the primordium cell layers that are derived from the mid-cortex of the primary root (Hossain et al., 2016; Shrestha et al., 2020; Xiao et al., 2014). Related to this, several lines of our investigation on the function of *LSH1/LSH2* during nodulation have pointed towards the hypothesis that *LSH1/LSH2* and *NF-YA1* may at least in part act within the same regulatory pathway to promote these processes. Consistent with this hypothesis we observed very similar phenotypes between *lsh1/lsh2* and *nf-ya1*, with an increased ratio of white to blue *pNifH-GUS* expressing nodules, an increase in nodule number (Figures 2A-B, S2H and S4A-D) and an increased ratio of aborted cortical infection threads (Figures 3A-B and S4E-F). Furthermore, we observed a reduction of cell divisions and cell layers in the *nf-ya1* nodule primordia similar to *lsh1/lsh2* (Figures 3C, S3B and S4G). However, there are also differences between *lsh1/lsh2* and *nf-ya1* nodules, especially in the overall nodule morphology and maintenance of *pNifH-GUS* expression (Figures S4A-C).

**Figure 5.**
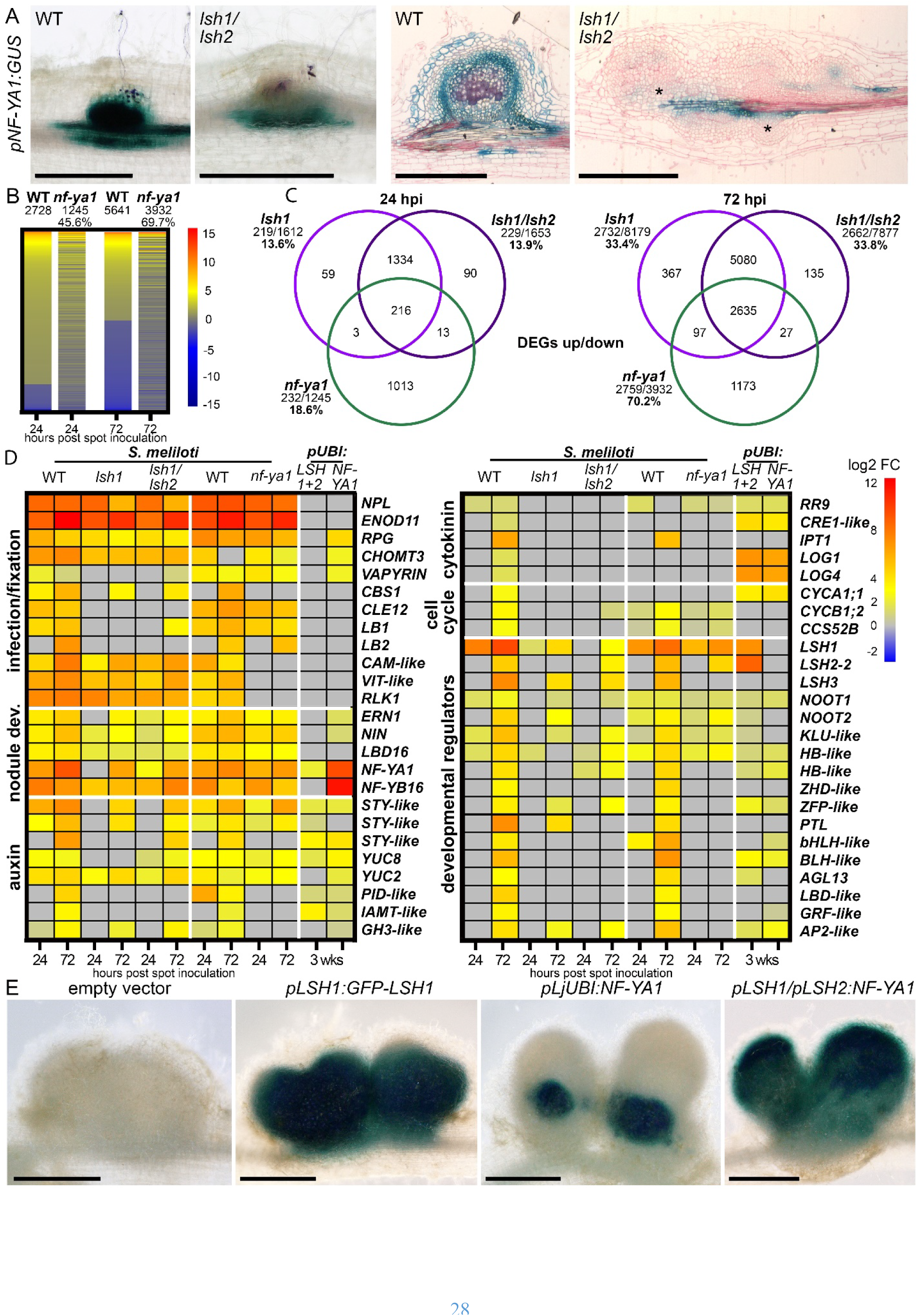
*LSH1/LSH2* partly function through the cortical activation of *NF-YA1*. (A) Expression pattern of *NF-YA1* in WT and *lsh1/lsh2* visualized by GUS staining (blue) in whole mount images (left) and nodule sections (right). Rhizobial expressed *LacZ* is stained magenta. Ruthenium Red demarks cell walls in sections. Black asterisks indicate vascular expression restricted at the nodule base. Scale bars: 500 µm. (B) Heatmaps of all differentially expressed genes in response to *S. meliloti* spot inoculation in WT and in *nf-ya1* at 24 and 72 hpi. Expression levels are depicted as log_2_ fold changes (log_2_ fold changes ≥ +/-1, p-value < 0.05). Comparison of DEGs as described in Figure 4A. Percentages indicate the proportion of DEGs in WT that are not expressed in the mutant and therefore dependent on *NF-YA1*. (C) Related to Figure 5B. Comparisons of all DEGs dependent on *lsh1* (light purple), *lsh1/lsh2* (dark purple) and *nf-ya1* (green) up and down regulated at 24 hpi and 72 hpi. Genes with log_2_ fold changes of ≥ +/-1, p- value < 0.05 were included in this analysis. (D) Heatmap of selected functional groups of DEGs in WT, *lsh1*, *lsh1/lsh2* and *nf-ya1* at 24 and 72 hours post *S. meliloti* spot inoculation and in response to combined ectopic expression of *LSH1/LSH2* (*pUBI:LSH1/LSH2*) or *NF-YA1* (*pLjUBI:NF-YA1*) compared to empty vector control in 3-week old WT (jemalong) hairy roots under non-symbiotic conditions. Fold changes compared to controls are depicted in log2 scale with the significance threshold of p-value < 0.05. Also see Data S1. (E) Whole mount images of nodules on hairy roots of *lsh1/lsh2* plants transformed with empty vector control*, pLSH1:LSH1*, *pLjUBI:NF-YA1,* and combined *pLSH1:NF-YA1/pLSH2:NF-YA1* at 28 dpi with *S. meliloti* expressing *pNifH:GUS*. GUS staining (blue) indicates the expression of the bacterial *pNifH* promoter (also see Table S2). Scale bars: 500 µm.

To further investigate the commonalities and differences in *LSH1/LSH2* and *NF-YA1* functions, we performed RNA-Seq on rhizobial spot inoculated *nf-ya1* and WT root sections at 24 and 72 hpi and compared the gene dependencies of rhizobial-induced genes between *NF-YA1* and *LSH1/LSH2* (Figures 5B-C and S5C, Data S1). *NF-YA1* controls a comparatively smaller subset of the rhizobial-induced genes than *LSH1/LSH2*: 46% and 70% of rhizobial-responsive genes were dependent on *NF-YA1* at 24 hpi and 72 hpi, respectively (Figure 5B): compare this to >90% for *LSH1/LSH2* (Figure 4A). While there was a surprisingly small overlap of < 20% between all *NF- YA1*-dependent and *LSH1/LSH2*-dependent genes at 24 hpi, the overlap of *NF-YA1*-dependent genes increased to 71% at 72 hpi, (Figures 5C and S5C). In addition, we constitutively expressed *NF-YA1* in hairy roots under non-symbiotic conditions (*LjUBI:NF-YA1*). Genes that showed strong transcriptional responses in the gain and loss of *LSH* and *NF-YA1* further highlight the role of local auxin biosynthesis, transport and conjugation, cytokinin signaling and cell cycle regulation during early nodulation, but also include several shoot-expressed growth regulators (Figure 5D and Data S1). Furthermore, this dataset confirmed that the expression of *LSH1/LSH2* is not dependent on *NF-YA1* (Figures 1B and 5D). Together this suggests that these regulators have independent, additive functions at the early nodule primordium stage and converge on similar regulatory pathways at the timepoint of early cortical infection and nodule differentiation. Based on this, we hypothesized that these regulators act within similar pathways and that the reduced cortical expression of *NF-YA1* might at least in part explain the reduced bacterial colonization and N-fixation phenotype observed in *lsh1/lsh2*. To test this we ectopically expressed *NF-YA1* under the constitutive *LjUBI* promoter or under the *pLSH1* and *pLSH2* promoters in *lsh1/lsh2* roots. Both modes of *NF-YA1* expression resulted in a partial rescue of *lsh1/lsh2,* leading to 25% of nodules with functional N-fixation, based on *pNifH-GUS* (Figure 5E, Table S2), revealing a functional link between *LSH1/LSH2* and *NF-YA1* during nodule organogenesis.

### *LSH1/LSH2* and *NOOT1/NOOT2* function in the same pathway during nodule organogenesis

Our RNA-Seq results also suggested a dependency of *NOOT1/NOOT2* expression on *LSH1/LSH2*. To validate this, we performed promoter GUS analysis in hairy roots expressing *pNOOT1:GUS- tNOOT1* and *pNOOT2:GUS-tNOOT2* in wild type and *lsh1/lsh2*. Both *NOOT* reporters showed expression in the inner tissue layers at the base of the developing nodule and in the nodule primordium in the wild type (Figure 6A), but in *lsh1/lsh2*, we observed a moderate reduction in expression of *NOOT1* and a loss of expression of *NOOT2* in nodule primordia (Figure 6A).

**Figure 6.**
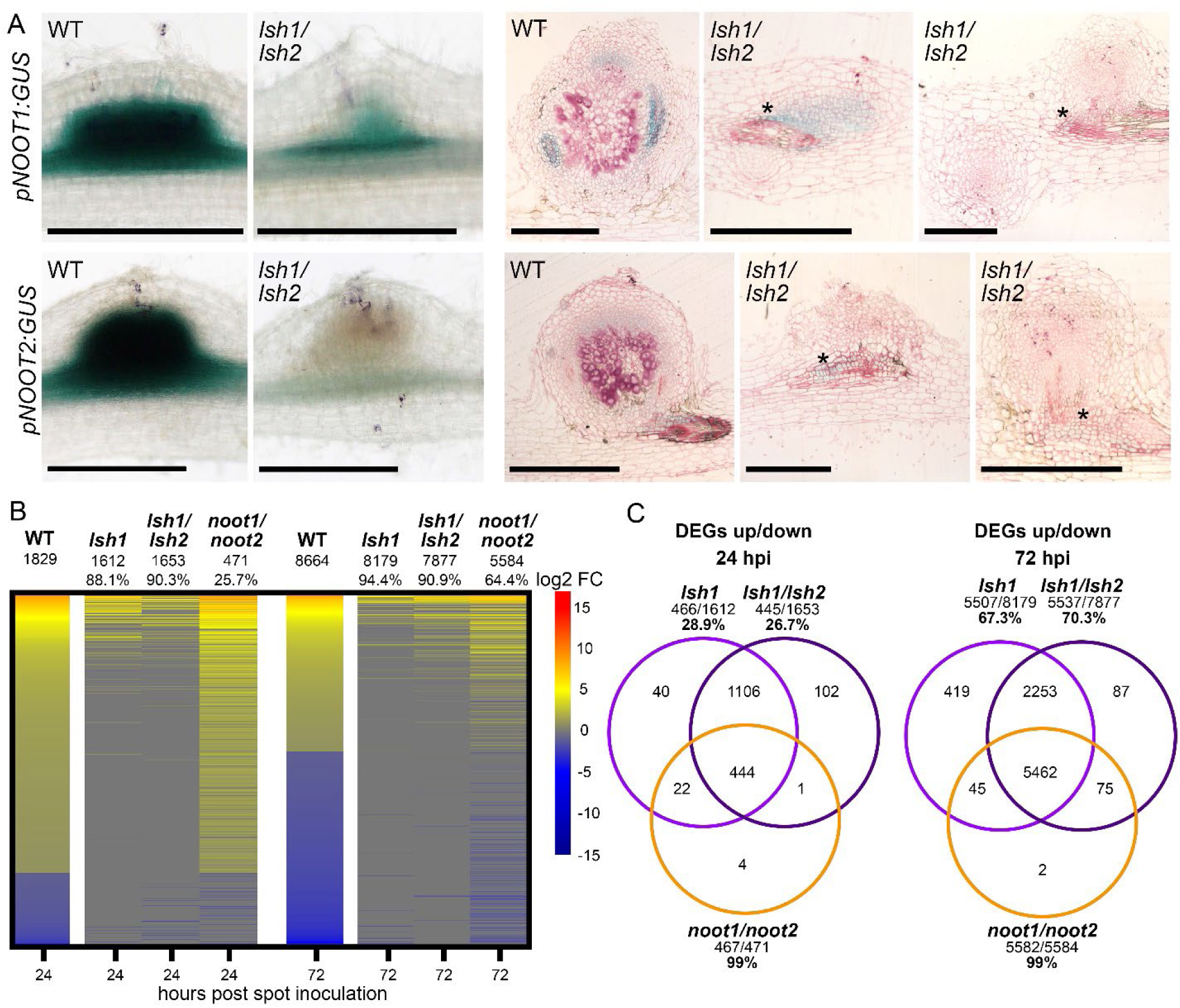
*LSH1/LSH2* promote the expression of and act together with *NOOT1/NOOT*2 in the same regulatory pathways. (A) Expression patterns of *NOOT1* and *NOOT2* in WT and *lsh1/lsh2* nodules, visualized by GUS staining (blue) in whole mount images (left) and nodule sections (right). Rhizobial expressed *LacZ* is stained magenta. Ruthenium Red demarks cell walls in sections. Black asterisks indicate vascular expression at the nodule base. Scale bars: 500 µm. (B) Related to Figures 4A-B and S5A-C, also see Data S1. Heatmaps of all DEGs in WT, *lsh1*, *lsh1/lsh2* and *noot1/noot2* at 24 and 72 hpi. Expression levels are depicted as log_2_ fold changes (log_2_ fold changes ≥ +/-1, p-value < 0.05). Comparison as described in Figure 4A. Percentages indicate the proportion of DEGs in WT that are not differentially expressed in the mutants and therefore dependent on *LSH1*, *LSH1/2* and *NOOT1/2*. (C) Comparisons of all DEGs dependent on *lsh1* (light purple), *lsh1/ lsh2* (dark purple) and *noot1/ noot2* (orange) up and down regulated at 24 hpi and 72 hpi. Genes with log_2_ fold changes of ≥ +/-1, p-value < 0.05 were included in this analysis.

It has previously been shown that several orthologs of the *BOP* (*NOOT*) genes function together with members of the *LSH* family to regulate organ development in the shoot (He et al., 2020; Xu et al., 2016). To test whether *LSH* and *NOOT* function together within the same regulatory pathway during symbiotic nodule development, we included *noot1/noot2* into our time-resolved expression and functional analyses. Unlike *lsh1/lsh2* nodule primordia which showed a clear reduction in the periclinal cell divisions of the root cortex, *noot1/noot2* primordia showed cell cycle activities comparable or greater than wild type (Figures 3C and 7B) and wild-type rhizobial infection, as previously reported (Magne et al., 2018). However, at later stages of nodule development, *noot1/noot2* showed similar defects to *lsh1/lsh2* in rhizobial colonization, resulting in a large proportion of partially or completely uncolonized nodules (Figures 3A-B and S6A-B). Loss of *NOOT1/NOOT2* affects a much smaller subset of the rhizobial-induced gene set than the loss of *LSH1/LSH2:* 25.75% and 64.45% of rhizobial-responsive genes were not differentially expressed in the *noot1/noot2* mutant at 24 hpi and 72 hpi, respectively (Figure 6B), compared to >90% in *lsh1/lsh2* (Figure 4A). There was an 99% overlap between the genes that were not responding to rhizobial inoculation in the *lsh1/lsh2* and in the *noot1/noot2* mutants, suggesting that the effect on gene expression caused by loss of *NOOT1/NOOT2* is completely embedded in the *LSH1/LSH2* function (Figure 6C). The constitutive expression of *NOOT1*/*NOOT2* (*pLjUBI:GFP- NOOT1 pLjUBI:GFP-NOOT2*) revealed a substantial (75%) overlap with genes induced by overexpression of *LSH1/LSH2* (Figures S5B-C). Genes that showed strong transcriptional responses in the gain and loss of *LSH* and *NOOT* further highlight the role of growth regulators with pleiotropic functions in shoot development, auxin and cytokinin signaling and the requirement for the repression of root meristem regulators such as *PLT1* and *PLT2* during nodulation (Figures S5A). *LSH1/LSH2* control the expression of *NOOT1/NOOT2* genes during nodulation, but *NOOT1/NOOT2* has no effect on *LSH1* or *LSH2* expression (Figures 4B and S5A). These studies suggest that *NOOT1/NOOT2* function downstream of *LSH1/LSH2* and the lack of their expression, at least in part explains the *lsh1/lsh2* phenotype, especially in the later stages of nodule development. Consistent with such a hypothesis we observed genetic interactions between *LSH* and *NOOT,* with a *lsh1/noot1* double mutant recapitulating the phenotype of a *lsh1/lsh2* double mutant (Figures 7A-B and S6A-C). A striking aspect of the *noot* mutants are the emergence of lateral roots from the tip of nodules (Magne et al., 2018). We observe this phenotype in *lsh1/lsh2*, but at a lower frequency to that observed *in noot1/noot2* (Figures 7A-B and S6C), consistent with the loss of repression of root meristem genes such as *PLT1/PLT2* in both double mutants (Figures 4B and S5A). The phenotypic resemblance between the *lsh1/noot1* and *lsh1/lsh2* mutants was also observed at earlier timepoints, where we observed early infection defects and a clear reduction in the periclinal cell divisions of cells derived from the root cortex in *lsh1/noot1* as initially observed in the *lsh1/lsh2* mutant (Figures 3C, 7C and S6A-B). We conclude that *LSH1* and *LSH2* control *NOOT1* and *NOOT2* and this regulation, in part explains the loss of function *lsh1/lsh2* phenotype.

**Figure 7.**
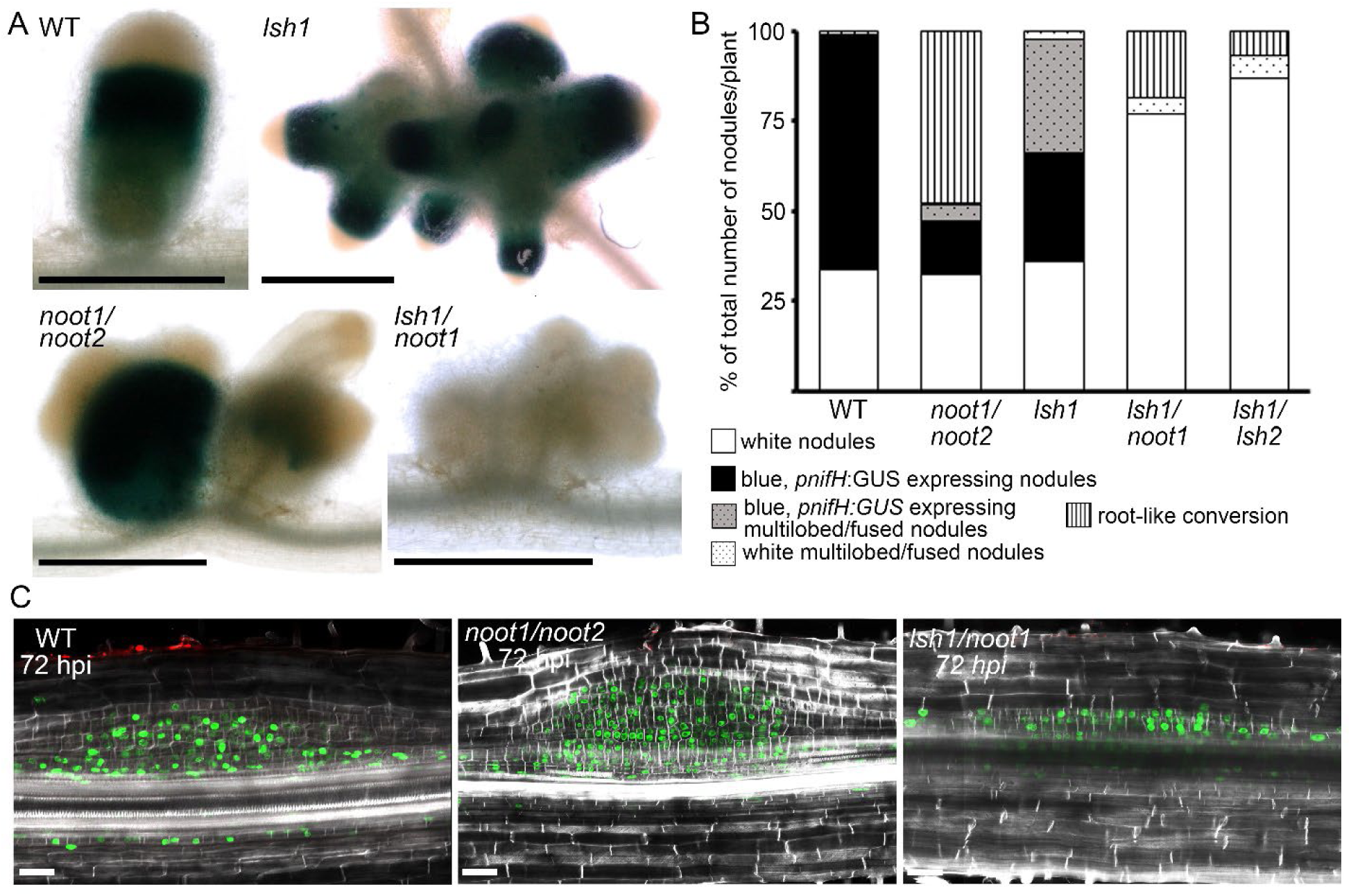
*LSH1/LSH2* and *NOOT1/NOOT*2 function synergistically to confer nodule organ identity. (A) Whole mount images of WT, *lsh1*, *noot1/noot2* and *lsh1/ noot1* nodules at 21 days post *S. meliloti* inoculation. GUS staining (blue) indicates the expression of the bacterial *pNifH.* Scale bars: 500 µm. (B) Distribution of different nodule morphologies and N-fixation (*pnifH:GUS* staining) at 21 dpi depicted as percentage of the total nodule number per plant, WT (n = 37), *noot1/noot2* (n= 44), *lsh1* (n= 33), *lsh1/noot1* (n = 35), *lsh1/lsh2* (n = 40) (also see Figure S6C). (C) Optical sections of WT, *noot1/noot2* and *lsh1/noot1* root sections 72 h post rhizobial spot-inoculation (n >15 per genotype). *Sm2011-mCherry* bacteria in red, cell walls in white (fluorescent brightener) and EdU-labelled nuclei indicating DNA replication in green. White arrowheads indicate periclinal cell divisions. Scale bars: 50 µm. For comparison to *lsh1/lsh2* also see Figures 3C and S3B).

## DISCUSSION

The initiation of symbiotic root nodules and lateral roots converges on the upregulation of the transcription factor *LBD16,* that allows the initiation of cell divisions in the inner root tissue layers, in part through the activation of local auxin biosynthesis (Schiessl et al., 2019; Soyano et al., 2019). Despite these similarities during the initiation stages, the development of nodules and lateral roots diverge, with nodules differentiating into organs that can intracellularly accommodate rhizobial bacteria and provide the environment for biological N-fixation. Here, we show that nodule-specific development is orchestrated by *LSH1/LSH2*, two members of the *LSH* transcription factor family with previously reported pleiotropic functions in shoot development. Like *LBD16*, *LSH1/LSH2* are controlled by *NIN* downstream of cytokinin signaling, suggesting parallel activation of two pre-existing conserved programs: a root-specific initiation program and a primordium identity program by *NIN* during the promotion of nodule organogenesis. Whereas *LBD16* is essential for the initiation of the nodule primordia (Schiessl et al., 2019; Soyano et al., 2019), *LSH1/LSH2* appear to be required for the specification of nodule primordia which is associated with cell divisions in the mid-cortex, responsible for the production of cells able to be intracellularly colonized by rhizobial bacteria. *LSH1/LSH2* function includes the cortex-specific promotion of the previously identified nodule regulators *NF-YA1* and *NOOT1/NOOT2* and our work positions *LSH1/LSH2* as integrators of nodule organ identity establishment and maintenance downstream of *NIN*. We propose, that whereas *NIN* promotion of *LBD16* initiates cell divisions in a manner parallel to lateral root development, *NIN* induction of *LSH1/LSH2* allows the emergence of nodule-specific features and functions.

*LSH1/LSH2* controls the expression of *NF-YA1*, in particular in the cortex-derived layers of the nodule primordium, positioning *LSH1/LSH2* as upstream regulators of nodule organ identity at the early stages of development. Previously, *NF-YA1* has been characterized as a direct downstream target of *NIN* and shown to be involved in nodule initiation and the progression of bacterial infection (Laporte et al., 2014; Soyano et al., 2013; Soyano et al., 2019). Based on our results, we propose that *LSH1/LSH2* promotes *NF-YA1* expression specifically in the cortical cell layers associated with the specification of the nodule primordium, perhaps subsequent and in parallel to its direct upregulation by *NIN* in the pericycle and endodermal cells at the base of the developing nodule. This is consistent with the modest overlap in *NF-YA1-* and *LSH1/LSH2-* dependent target genes at early stages of development, but the strong overlap in gene expression dependencies at later stages of nodule differentiation. Functional overlaps between *LSH1/LSH2* and *NF-YA1* are particularly pronounced in the regulation of genes involved in local auxin biosynthesis, polar auxin transport and cytokinin biosynthesis and signaling. Cross talk between polar auxin transport and cytokinin signaling has previously been shown to determine cell division patterns and the onset of cell differentiation in the transition zone of the root and during lateral root development (Ioio et al., 2008; Laplaze et al., 2007; Marhavý et al., 2011; Marhavý et al., 2014). Equally, the maintenance of cytokinin homeostasis and cytokinin signaling have been shown to play vital roles in promoting nodule organogenesis and endosymbiotic host cell specification in legumes (Mortier et al., 2014; Reid et al., 2015; Tan et al., 2020)

In contrast to *NF-YA1, NOOT1/NOOT2* function appears to be entirely controlled by *LSH1/LSH2*, with very severe reductions in *NOOT1* and *NOOT2* expression in *lsh1/lsh2* and an almost complete overlap in the effect on rhizobial-induced differential gene expression between the *noot1/noot2* and *lsh1/lsh2* mutants. The severe nodulation defects caused by the combined loss of *LSH1* and *NOOT1*, which are comparable to the defects found in the *lsh1/lsh2* double mutant suggests a supportive role of the *NOOT* genes for *LSH1/LSH2* function. Together with previous studies in which members of the *BOP* family support the function of members of the *LSH* gene family post-translationally, we propose a regulatory feedback loop in which *LSH* promotes the expression of their synergistic interaction partners during nodulation (He et al. 2020, Xu et al. 2016). Through the combined control of the early *NF-YA1* and the late *NOOT1/NOOT2* function, *LSH1/LSH2* appear to promote specific cell divisions in middle cortical cell layers associated with the establishment and maintenance of nodule organ identity throughout nodule development and the transition to the N-fixing state. We propose a model for nodule formation: NIN initiates the nodule primordium through induction of *LBD16*, which in turn activates local auxin biosynthesis to initiate cell divisions; in parallel NIN activates the expression of *LSH1/LSH2,* that control and maintain nodule organ identity, in part through the regulation of *NF-YA1* and *NOOT1/NOOT2* creating cells able to accommodate N-fixing bacteria; a final step in the maturation of the nodule is controlled by NIN processing that allows direct NIN regulation of late nodule-associated genes involved in supporting N-fixation (Feng et al., 2021; Schiessl et al., 2019).

Loss of both *LSH1/LSH2* and *NF-YA1* have strong effects on the progression of the infection thread into the root cortex. It remains to be elucidated how *LSH1/LSH2* that are restricted in their expression to the inner tissue layers of the developing nodule primordium can have an equally severe effect on infection thread progression as *NF-YA1,* which is expressed in the epidermis (Laporte et al., 2014). Recent work by Goto et al. (2022) implicated methylated auxin (MeIAA) in the promotion of the epidermal infection process in *L. japonicus.* MeIAA is a non-polar auxin conjugate that can move without the need of active auxin transport (Goto et al., 2022; Qin et al., 2005). The enzyme required for its biosynthesis, *IAA CARBOXYL METHYLTRANSFERASE* (*IAMT1*), is dependent on *LSH1/LSH2* and *NF-YA1* function. MeIAA is a potential candidate for the yet to be discovered signal that crosses tissue layers to coordinate the progression of the epidermal infection thread with the cortical organogenesis process. The absence of this potential signal may explain the infection defects in *LSH1/LSH2*.

In the shoot, *LSH* genes have been shown to regulate the activity of meristems and the growth of lateral organs with a proposed function in promoting the appropriate amount of undifferentiated cells that can serve as a canvas for the differentiation processes that result in complex organs with intricate features and functions (He et al., 2020; MacAlister et al., 2012; Yoshida et al., 2013). This is exemplified by their role in the development of indeterminate compound inflorescences in tomato and rice (MacAlister et al., 2012; Yoshida et al., 2013) and the development of stipules with complex serrations in *M. truncatula,* as demonstrated here. We propose that these previously described shoot functions of *LSH* and its co-recruited shoot-expressed regulatory subnetwork are of relevance during nodule development where early cell patterning processes need to be spatially and temporally coordinated with the arrival of the colonizing bacteria. This recruitment appears to be specific to nodules as we observe no induction of *LSH1* or *LSH2* during early lateral root development (Schiessl et al., 2019), suggesting that symbiotic root nodule evolution involved the recruitment of a pre-existing program with pleiotropic functions in the shoot, into the symbiotic program to facilitate the unique morphological and functional features of the symbiotic root nodule. This represents a key point of divergence between nodule and lateral root development.

The neo-functionalization of the nodule-specific transcription factor *NIN* and the associated evolution of cis-regulatory DNA binding sites in the promoter regions of its downstream targets led to the recruitment of a lateral root-like organ initiation program into the symbiotic interaction with rhizobial bacteria (Dong et al., 2021; Griesmann et al., 2018; Schiessl et al., 2019; Soyano et al., 2019). Similarly, it can be hypothesized that the neofunctionalization of *NIN* provided the opportunity to recruit a growth regulatory network with pleiotropic functions in the shoot into the N-symbiotic root context, thereby promoting the expansion and diversification of the regulatory function of, for example, *LSH1/LSH2* and their associated pre-existing downstream regulatory subnetworks to the legume root. This notion is in line with the common principle of morphological evolution as proposed by Caroll et al. (2008) in which changes in the spatial and temporal gene expression of pre-existing developmental regulators and their associated downstream networks, lead to trait divergence and the diversification of novel organ forms and functions (Carroll, 2008). The parallel recruitment of a root initiation program and primordium identity program from the shoot that dictate nodule form and function are essential in non-legumes species that are targets for engineering N-fixation.

## ACKNOWLEDGEMENTS

We thank Raymond Wightman, Gareth Evans, Kim Findlay, and Elaine Barclay for support with microscopy, Anne Edwards for help with the acetylene reduction assay and Mandana Miri for critical comments on the manuscript. We thank Pascal Ratet for providing *noot1/noot2* mutant seeds. This work was supported by the Bill and Melinda Gates Foundation and the UK Foreign, Commonwealth and Development Office (OPP1172165) through Engineering the Nitrogen Symbiosis for Africa (ENSA) project and the Gatsby Foundation (GAT3395/GLH).

## AUTHOR CONTRIBUTIONS

K.S. designed and performed the experiments and analyzed the data with assistance from M.O. and N.S.; T.L. analyzed RNA sequencing (RNA-Seq) data; P.C.B. performed the phylogenetic analysis; M.B-B. provided resources and performed the acetylene reduction assay; K.S.M. and J.W. provided *Medicago* mutants; K.S. and G.E.D.O supervised the work and wrote the manuscript.

## STAR METHODS

### LEAD CONTACT AND MATERIALS AVAILABILITY

Further information and requests for resources and reagents should be directed to and will be fulfilled by the lead contact, Giles Oldroyd, subject to material transfer agreements. This study did not generate unique reagents.

### EXPERIMENTAL MODEL AND SUBJECT DETAILS

#### Plant material and *Sinorhizobium meliloti* strains including growth conditions

*M. truncatula* ecotypes jemalong, cultivar Jester, and ecotype R108 were used in this study. Jemalong was used to perform hairy root transformations and as wild type for comparisons to *cre-1*, *nin-1*, and *nfy-a1-1*, previously described (Marsh et al., 2007; Plet et al., 2011). All *Tnt1* retrotransposon insertion lines described in this manuscript (*NF17203 (lsh1-1)*, *NF1304 (lsh1-2)* and *NF14992 (lsh2-1*) were derivatives of the R108 ecotype and obtained from the *Tnt1* Retrotransposon Mutant Collection (Oklahoma State University, Stillwater, USA; previously Noble Research Institute, LLC., Ardmore USA; (Cheng et al., 2014). Previously described *NF2717* (*noot1-1*), *NF5464* (*noot2-1)* were a gift from Pascal Ratet, Institut des Sciences des Plantes de Paris Saclay, Orsay, France. As such R108 was used as the wild type for analysis of these mutants. Genotyping was performed using Tnt1-F and Tnt1-R oligos combined with the corresponding forward and reverse oligos encompassing the insertions (Table S1A).

Rhizobial bacterial strains of *Sinorhizobium meliloti* 2011 expressing *pXLGD4 (hemA: lacZ), pNifH:GUS,* or *mCherry* were used in this study (Lerouge et al., 1990; Starker et al., 2006).

Seeds were scarified, surface sterilized with 10 % (v/v) bleach solution, stratified for 3 days at 4 °C and germinated on water agar plates. Plants were grown in sterile conditions in controlled environment rooms at 22 °C (80 % humidity, 16 h light/8 h dark, 300 μmol m2 s-1 light intensity) on filter paper-lined agar media in sealed plates unless otherwise specified. For rhizobial spot inoculation seedlings were grown for 2 days on buffered nodulation medium (BNM) (Ehrhardt et al., 1992) supplemented with 1 µM aminoethoxyvinylglycine (AVG; Sigma-Aldrich Company Ltd, Darmstadt, Germany) at 22 °C (16 hours light/8 hours dark, 300 μmol m-2 s-1 light intensity). *Sinorhizobium meliloti* strain 2011 (Lerouge et al., 1990) was grown in minimal medium supplemented with 3 µM luteolin (Sigma-Aldrich Company Ltd, Darmstadt, Germany) and diluted to a final concentration of 0.02 OD 600 nm using Fahraeus medium (Boisson-Dernier et al., 2001). The mock treatment consisted of Fahraeus medium with luteolin diluted to an equivalent concentration as the inoculum. Approximately 1 µL of *S. meliloti* suspension or mock treatment was inoculated onto the susceptibility zone (where the root hairs first appear) and marked by puncturing the filter paper alongside the site of inoculation. After 24 and 72 hours, 2 mm sections of the root alongside the site of inoculation were harvested for RNA isolation or microscopy.

For spray inoculation of *S. meliloti* for plants grown on plates, seedlings were grown under similar conditions as described above. Roots of 1-day-old seedlings were covered with filter paper and sprayed with 2 ml *S. meliloti* of final concentration 0.02 OD 600 nm grown in minimal medium without luteolin. For inoculation of hairy roots and for phenotyping assays of mature nodules, plants were transferred to terragreen:sharp sand mix (1:1) (Oil-DriCompany, Wisbech, UK) in P40 trays and left to grow for 7 days before inoculation with *S. meliloti 2011* (2 mL of overnight culture per plant diluted in liquid BNM to 0.02 OD 600 nm). Plants were grown for up to a further 4 weeks for nodule quantification and histochemical staining.

#### Bacterial strains

*Agrobacterium rhizogenes* strain *AR1193* was used to introduce all binary vectors used in this study (*DsRed* as transformation marker) to *M. truncatula* jemalong seedlings following a previously published transient hairy root transformation protocol (Boisson-Dernier et al., 2001).

### METHOD DETAILS

#### Construct production

The Golden Gate modular cloning system was used to prepare the plasmids (Weber et al., 2011). All Level 0s used in this study are held for distribution in the ENSA project core collection (https://www.ensa.ac.uk/) and are listed along with the binary plasmid details in Table S1A. Sequences were domesticated, synthesized and cloned into pMS (GeneArt, Thermo Fisher Scientific, Waltham, USA). Sequence information for *LSH1* (*Medtr1g069825), LSH2 (Medtr7g097030), NF-YA1 (Medtr1g056530), NOOT1 (Medtr7g090020)* and *NOOT2 (Medtr1g051025)* were obtained from the *M. truncatula* Mt4.0v1 genome via Phytozome (https://phytozome.jgi.doe.gov) (Rokhsar et al., 2011).

#### Hormone and chemical treatments

AVG and 6-Benzylaminopurine (BAP; Sigma-Aldrich Company Ltd, Darmstadt, Germany) were dissolved in water. Mock treatments were equal volumes of each solvent in the agar media. For BAP plate treatments (100 nM) 2-day old seedlings were grown on BNM plates for 24 hours with either BAP or mock and supplemented with 1 µM AVG to replicate spot inoculation conditions.

#### Gene expression analysis

For rhizobial spot inoculation time-course experiments, roots were dissected as 2- to 3-mm segments around the spot of inoculation or mock treatment. For BAP response experiments, segments were dissected around the susceptibility zone marked at the time of treatment. About 50 to 60 segments were pooled to obtain 1 biological replicate, with 3-6 biological replicates per treatment/genotype were analyzed. RNA was extracted using the RNeasy Micro Kit (Qiagen, Germantown, USA) and the RNase free DNase kit (Qiagen, Germantown, USA) was used to remove genomic DNA. For reverse transcription of 1 µg total RNA, Transcriptor First Strand cDNA Synthesis Kit was used according to the manufacturer’s instructions (Roche Diagnostics GmbH). Quantitative real-time polymerase chain reactions (qRT-PCR) were performed in technical triplicates in the LightCycler 480 System using LightCycler 480 SYBR green I master (04707516001, Roche Diagnostics GmbH, Mannheim, Germany) in a total reaction volume of 10 μl. The primer pairs used for gene expression analysis are listed in Table S1A.

#### RNA-Seq

RNA sequencing (RNA-Seq) was performed by Novogene Europe (Cambridge, UK). RNA-Seq libraries were prepared with the Illumina TruSeq® Stranded mRNA HT kit The sequencing of the libraries was performed on the Illumina NovaSeq 6000 next generation sequencing system with the read length of 150 bp paired-end reads resulting in 20 million reads per sample.

#### Assessment of shoot-related phenotypes

Flowers and stipules were dissected from the main shoot, arranged on double-sided tape and directly imaged using the Keyence VHX-5000 microscope equipped with a range of zoom lenses from 20x to 1000x (Keyence Ltd, Milton Keynes, UK).

#### Assessment of lateral root phenotypes

Seedlings were grown on modified Fahraeus medium (Boisson-Dernier et al., 2001) plates for 14 days before lateral root number and length was scored.

#### Acetylene reduction assay

Nitrogenase activity was measured *in vivo* in mature nodules of terragreen:sand grown plants at 21 days post inoculation with *S. meliloti 2011* by the acetylene reduction assay (Dilworth, 1966; Schöllhorn and Burris, 1967). Freshly harvested nodules were immediately transferred to BD Vacutainer tubes (3ml, 13×75mm, Thermo Fisher Scientific, Waltham, USA) prior to the injection of 10% (v/v) acetylene (C2H2). After 1 hour incubation, ethylene (C2H4) was quantified using a Perkin Elmer Clarus 480 gas chromatograph equipped with a HayeSep® N (80-100 MESH) column. The injector and oven temperatures were kept at 100 °C, while the FID detector was set at 150 °C. The carrier gas (nitrogen) flow was set at 8 - 10 mL/min. An ethylene calibration curve was prepared from chemical decomposition of ethephon (Sigma C0143) in 10 mM Na2HPO4 pH 10.7 as described previously (Zhang and Wen, 2010) . Nitrogenase activity is reported as nmol of C2H4 per nodule per hour.

#### Histochemical assays and cellular stains

For GUS staining roots were washed in water and immediately fixed in 90 % acetone on ice for 1 hr. Subsequently, the acetone was replaced by a wash solution containing 50 mM phosphate buffer pH 7.2. The wash buffer was replaced by GUS staining buffer containing 50 mM phosphate buffer pH 7.2, 0.5 mM K3Fe(CN)6 (potassium ferricyanide), 0.5 mM K4Fe(CN)6 (potassium ferrocyanide) and 2 mM 5-bromo-4-chloro-3-indolyl-beta-D-glucuronide (X-Gluc, Melford Laboratories Ltd., Ipswich, UK), vacuum infiltrated for 15 min and incubated at 37°C for 6 - 12 hrs. For X-Gal staining, the tissue was washed in 50 mM phosphate buffer pH 7.2 and fixed in 2.5 % glutaraldehyde by vacuum infiltration for 15 min and incubation at room temperature for 1 hr. Tissue was washed 3X in Z-buffer containing 100 mM phosphate buffer pH 7, 10 mM KCl, 1 mM MgCl2 and incubated in X-Gal staining buffer (Z-buffer supplemented with 5 mM K3Fe(CN)6, 5 mM K4Fe(CN)6 and 0.08% Magenta-5-Bromo-6-chloro-3-indolyl-B-D-galactopyranoside; (X-Gal, Melford Laboratories Ltd., Ipswich, UK) or 5-Bromo-4-chloro-3-indolyl β-D-galactopyranoside (blue X-Gal, Sigma-Aldrich Company Ltd, Darmstadt Germany) at 28°C for 6-12 hrs and washed with water 3X. Subsequently, nodules and root tissue were dehydrated in an ethanol series and stored in 70% EtOH at 4°C. For imaging of whole mount tissue, stained samples were mounted on glass slides in 70% EtOH and imaged using the Keyence VHX-5000 microscope equipped with a range of zoom lenses from 20x to 1000x (Keyence Ltd, Milton Keynes, UK).

#### Tissue Sections

Root segments and nodules were fixed with 4% formaldehyde (Sigma-Aldrich Company Ltd, Darmstadt Germany) in 1X PBS (10X PBS contains 1.37 M NaCl, 27 mM KCl, 100 mM Na2HPO4, and 18 mM KH2PO4) with vacuum infiltration for 15 min and at 4°C overnight. The fixed material was dehydrated in an ethanol series and subsequently embedded in Technovit 7100 (Kulzer Technik, Wehrheim, Germany) according to the manufacturer’s protocol. Embedded tissue was sectioned (8-10 µm) using a Leica microtome (Leica, Milton Keynes, UK), mounted on glass slides, stained in 0.1% Ruthenium Red (Sigma-Aldrich Company Ltd, Darmstadt Germany) dissolved in distilled water for 15 min and rinsed. Sections were imaged using a Zeiss Axioimager.M2 light microscope (Carl Zeiss AG, Oberkochen, Germany).

Combined 5-ethynyl-2-deoxyuridine (EdU; Invitrogen, Thermo Fisher Scientific, Waltham, USA) and modified pseudo-Schiff-propidium iodide (PI; Sigma-Aldrich Company Ltd, Darmstadt, Germany) staining, clearing and imaging was performed as previously published (Schiessl et al., 2019). In addition, staining with 5-ethynyl-2-deoxyuridine (EdU; Invitrogen, Thermo Fisher Scientific, Waltham, USA) was combined with the cell wall stain Calcofluor white - Fluorescent Brightener 28 disodium salt solution (Sigma-Aldrich Company Ltd, Darmstadt, Germany). For this, roots were transferred to growth medium supplemented with 10 µM EdU and grown for 4 additional hours as previously described (Schiessl et al., 2019). Subsequently, 1 cm root sections centred around the susceptibility zone were dissected and fixed in 4% formaldehyde in 1X PBS with vacuum infiltration for 15 min and incubation for 1 hr at room temperature. After 3 washes in 1X PBS, the root sections were incubated in solution containing 10 mM Alexa 488- azide (Invitrogen, Thermo Fisher Scientific, Waltham, USA) and 100 mM Tris pH 8.5 for 1 hr, followed by 30 min in solution containing 10 mM Alexa 488-azide, 100 mM Tris, 1 mM CuSO4, 100 mM ascorbic acid, pH 8.5. The roots were subsequently washed in water 3X and transferred to ClearSee solution (Ursache et al., 2018) for 24 – 72 hrs. For cell wall staining, root sections were transferred to ClearSee solution supplemented with 0.1% Fluorescent Brightener 28 disodium salt solution and incubated for 30 min (Ursache et al., 2018). For imaging root sections were mounted on glass slides in ClearSee solution and imaging was performed with a Zeiss 700 confocal scanning microscope using a 20X air lens objective with excitation and emission filters set to 405 nm/410–485 nm for Fluorescent Brightener 28 staining, at 488 nm/505–600 nm for EdU, and 488 nm/600-680 nm for rhizobial expressed *mCherry*, 488 nm/572–625 nm for propidium iodide (Carl Zeiss AG, Oberkochen, Germany). Images were processed using Zeiss software and FIJI (Schindelin et al., 2012).

#### Transmission electron microscopy analysis

Mature nodules of plate grown plants at 14 days post inoculation with *S. meliloti 2011* were cut from roots and immediately placed in a solution of 2.5% (v/v) glutaraldehyde in 0.05 M sodium cacodylate, pH 7.3 for fixation, and left overnight at room temperature. Samples were then placed in baskets and loaded into the Leica EM TP embedding machine (Leica, Milton Keynes, UK) using the following protocol. The fixative was washed out by three successive 15-minute washes in 0.05 M sodium cacodylate and post-fixed in 1% (w/v) OsO4 in 0.05 M sodium cacodylate for two hours at room temperature. The osmium fixation was followed by three, 15-minute washes in distilled water before beginning the ethanol dehydration series (30%, 50%, 70%, 95% and two changes of 100% ethanol, each for an hour). Once dehydrated, the samples were gradually infiltrated with LR White resin (London Resin Company, Reading, Berkshire, UK) by successive changes of resin:ethanol mixes at room temperature (1:1 for 1hr, 2:1 for 1hr, 3:1 for 1hr, 100% resin for 1 hr then 100% resin for 16 hrs and a fresh change again for a further 8 hrs) then the samples were transferred into gelatin capsules full of fresh LR White and placed at 60°C for 16 hrs to polymerize.

The material was sectioned with a diamond knife using a Leica UC6 ultramicrotome (Leica, Milton Keynes, UK) and ultrathin sections of approximately 90 nm were picked up onto 200 mesh copper grids which are formvar and carbon coated (EM Resolutions, Sheffield, England). The sections were stained with 2% (w/v) uranyl acetate for 1hr and 1% (w/v) lead citrate for 1 minute, washed in distilled water and air dried. The grids were viewed in a FEI Talos 200C transmission electron microscope (FEI UK Ltd, Cambridge, UK) at 200kV and imaged using a Gatan OneView 4K x 4K digital camera (Gatan, Cambridge, UK) to record DM4 files. For overview images, sections were stained in 0.05% Toluidine Blue O for 5 min and imaged using a Leica DM6000 compound microscope 20X air objective with bright field settings (Leica Microsystems, Wetzlar, Germany).

#### Phylogenetic analysis

LIGHT SENSITIVE SHORT HYPOCOTYL (LSH) proteins were detected in the *Arabidopsis thaliana*, *Medicago truncatula*, *Lotus japonicus*, *Solanum lycopersicum* and *Pisum sativum* proteomes using HMMER3.1b2 HMMSEARCH (hmmer.org). The inputs to this program were the Pfam hidden Markov model (HMM), PF04852 (Pfam release 30), and the protein data sets from the genome annotations of *A. thaliana* (Araport11), *M. truncatula* (Phytozome, V10), *L. japonicus* (http://www.kazusa.or.jp/lotus/, build 3.0), *S. lycopersicum* (Phytozome, ITAG3.2) and *P. sativum* (https://urgi.versailles.inra.fr/Species/Pisum, v1a). The protein sequences detected were aligned back to the HMM using HMMER3.1b2 HMMALIGN. Gap columns in the alignment that were not part of the HMM were removed, sequences with less than 70 % coverage across the alignment were removed and the longest sequence for each gene from the set of splice versions was used for phylogenetic analysis. Phylogenetic analysis was carried out using the MPI version of RAxML v8.2.9 (Stamatakis, 2014) with the following method parameters set: -f a, -x 12345, - p 12345, -# 100, -m PROTCATJTT. The tree was mid-point rooted and visualized using the Interactive Tree of Life (iToL) tool (Letunic and Bork, 2016).

### QUANTIFICATION AND STATISTICAL ANALYSIS

#### RNA-seq

Reads from the RNA-sequencing experiments provided as raw fastq data were quality controlled and mapped to the *M. truncatula* reference genome version 4.0 (Mt4.0v1) (Tang et al., 2014) using STAR (Dobin et al., 2013). The raw counts of aligned reads were calculated with featureCounts in R package Rsubread (Smyth et al., 2013). Non-metric multidimensional scaling was exploited to account for outliers. At least 3 biological replicates were always included in the full analysis. Genes that showed low expression throughout all samples were removed by measuring CPM (counts per million) values using R package edgeR (Robinson et al., 2010). Differentially expressed genes (DEGs) were identified by pairwise comparisons of raw counts of mock treatment versus experimental treatment, using the R package DESeq2 (Love et al., 2014) with the threshold of absolute fold change of over 2 and a false discovery rate (FDR) corrected p-value significance of 0.05. The heatmaps of differential expression were plotted with R package pheatmap. The synonyms for *M. truncatula* locus ids were obtained by manual curation and by retrieving names from the Uniprot database using BLAST matches (Altschul et al., 1990; Consortium, 2018). The descriptions for each gene were obtained from the Phytozome database.

#### qRTPCR

Expression values of minimum three biological replicates in three technical replicates were analyzed using the Pfaffl method with *histone H3* (*HH3*) as reference (Pfaffl, 2001). Statistical comparison was performed between WT and mutants or treatment and corresponding mock. Values depicted in bar charts are the mean of minimum 3 biological replicates ± SEM (Student’s t-test; * P < 0.05; ** P < 0.01, *** P < 0.001).

#### Phenotyping

Data on the morphology of nodule primordia, rhizobial infection threads and mature nodules was depicted in bar charts and in boxplots representing the percentage of total nodule number. Total number of serrations per stipules, total number of nodules per gram root tissue and acetylene reduction rates were depicted as boxplots. Boxplots show the median (thick line), second to third quartiles (box), minimum and maximum ranges (lines), and outliers (single points). Normal distribution of data was tested using the Shapiro-Wilk normality test. For pairwise comparisons statistical analysis was performed using either unpaired Student’s t-test, Wilcoxon test or Fisher’s exact test. For multiple comparisons, one-way analysis of variance (one-way ANOVA) or one-way Kruskal-Wallis rank sum test, followed by Tukey multiple comparisons of means or Dunn test. The R statistical package was used for these analyses. Samples size n is provided in the figure legends and refers to the number of individual plants unless indicated otherwise. Statistical tests and significance levels are provided in the figure legends.

### DATA AND CODE AVAILABILITY

The short-read sequencing data generated in this study have been deposited at the National Center for Biotechnology Information Gene Expression Omnibus, with accession number ---. Lists of differentially expressed genes are compiled in Data S1.

## SUPPLEMENTAL INFORMATION

Document S1 includes Figure S1-S6 and Table S1-S4.

