## Supplemental Figures S1-6, Tables S1-4 for "Light sensitive short hypocotyl (LSH) confer symbiotic nodule identity in the legume Medicago truncatula"

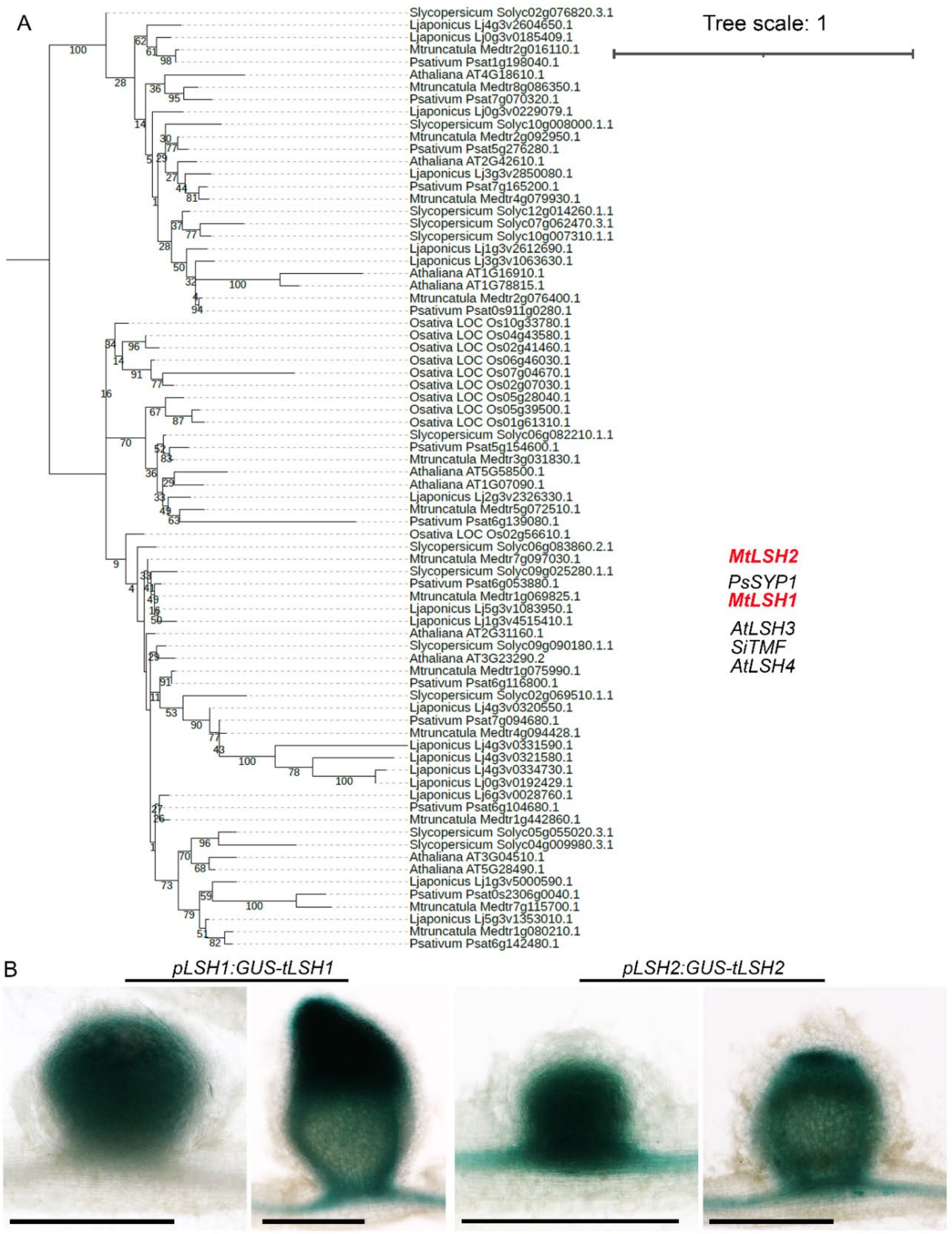


**Figure S1. Identification of *LIGHT SENSITIVE SHORT HYPOCOTYL (LSH)* genes expressed during nodule organogenesis in *Medicago truncatula*. Related to Figure 1.** (A) Phylogenetic analysis of the *LSH* gene family. The tree shows 76 genes from *Arabidopsis thaliana*, *Solanum lycopersicum*, *Lotus japonicus*, *Pisum sativum*, *Medicago truncatula* and *Oryza sativa*. Genes functionally characterized in this study are labelled in red. Tree scale: 1. (B) Expression patterns of *LSH1 and LSH2* during nodule development visualized by GUS staining (blue). Scale bars: 500 µm.

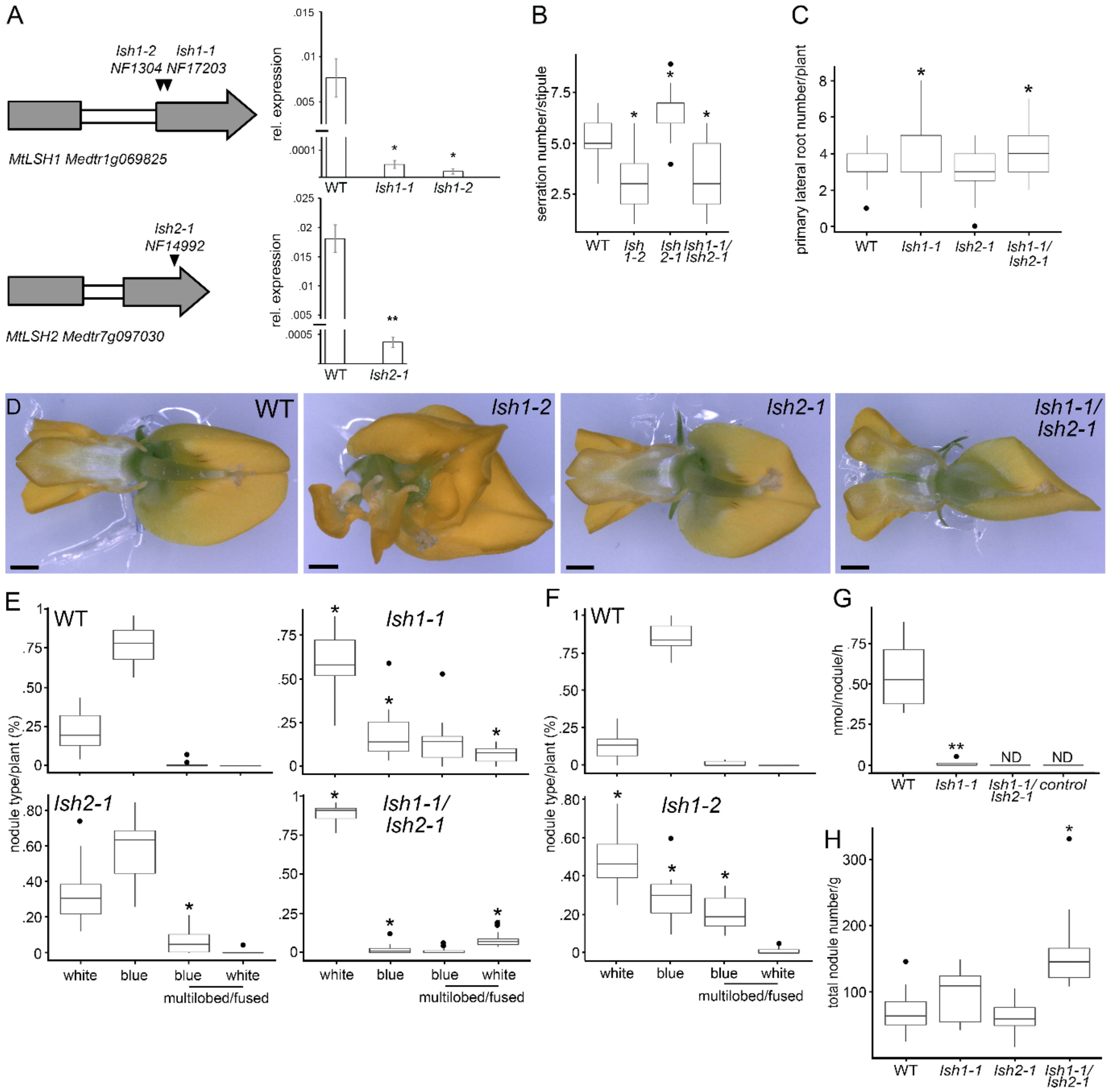


**Figure S2. *LSH1* and *LSH2* are required for the development of nitrogen-fixing nodules. Related to Figure 2.** (A) Location of exonic *Tnt1* retrotransposon insertions and comparison of transcript levels in WT and the corresponding exonic *Tnt1* retrotransposon insertion lines measured in 8-day old seedling roots, 7 days post spray inoculation with *S. meliloti*. Expression levels were measured by qRT-PCR and normalized to *HH3*. Statistical comparison was performed between WT and the insertion lines. Values are the mean of 3 biological replicates (>5 roots/replicate) ± SEM (Student's t-test; * $P < 0.05$; ** $P < 0.01$, *** $P < 0.001$). (B) Distribution of serration number per stipule collected from the third and fourth internode below the shoot apex (as depicted in Figure 2B) in WT (n = 22), *lsh1-2* (n = 16), *lsh2-1* (n = 15), *lsh1-1/lsh2-1* (n = 20). A one-way Kruskal-Wallis rank sum test showed that serration number is dependent on genotype (KW = 77.471, df = 3, p < 2.2e-16). Asterisks indicate significantly different means for *lsh1-2*, *lsh1-1/lsh2-1* (lower) and *lsh2-1* (higher) compared with WT, Dunn Test (95% confidence). (C) Distribution of primary lateral root number assessed in plate-grown 14-day old seedlings of WT (n = 49), *lsh1-1* (n = 40), *lsh2-1* (n = 47), *lsh1-1/lsh2-1* (n = 55). A one-way Kruskal-Wallis rank sum test showed that primary lateral root number is dependent on genotype (KW = 28.308, df = 3, $p < 3.13e-06$). Asterisks indicate significantly different means for *lsh1-1*, *lsh1-1/lsh2-1* compared with WT, Dunn Test (95% confidence). (D) Images of WT, *lsh1-2*, *lsh2-1* and *lsh1-1/lsh2-1* flowers as used for the phenotyping in Table S1. Scale bars: 500 µm. (E) Related to Figure 2C-E. Distribution of nodule morphologies/types categorised in "white", *pNifH:GUS* expressing "blue", *pNifH:GUS* expressing "blue – multilobed and/or fused" and "white multilobed and/or fused" per plant in percentage for WT (n = 15, boxplot top left), *lsh1-1* (n = 15, boxplot top right), *lsh2-1* (n = 14, boxplot bottom left), and *lsh1-1/lsh2-1* (n = 13, boxplot bottom right) at 28 days post *S. meliloti* spray inoculation. One-way Kruskal-Wallis rank sum tests showed that the distribution of nodule types is dependent on genotype (KW = 40.65 , df = 3, p = 7.759e-09 (white), KW = 47.00, df = 3, p = 3.468e-10 (blue), KW =26.181 , df = 3, p = 8.742e-06 (blue- multilobed and/or fused) and KW = 41.64, df = 3, p = 4.783e-09 (white multilobed and/or fused). Asterisks indicate significantly different means for *lsh1-1, lsh2-1* and *lsh1-1/lsh2-1* compared with WT, Dunn Test (95 % confidence). (F) Distribution of nodule morphologies per plant in percentage for WT (n = 13) and *lsh1-2* (n= 11) at 28 days post *S. meliloti* spray inoculation as described in (E). Student's t-tests showed that the distribution of the different nodule types is dependent on genotype; Asterisks indicated statistical significance *, $P < 0.05$; **, $P < 0.01$, ***, $P < 0.001$). (G) Comparison of *in vivo* nitrogenase activity in nodules of WT, *lsh1-1* and *lsh1-1/lsh1-2*, collected at 21 days post spray inoculation with *S. meliloti*. Nitrogenase activity was measured by the acetylene reduction assay and reported as nmol of $C_2H_4$ per nodule per hour. Nitrogenase activity levels were below detection threshold in *lsh1-1/lsh2-1* nodules. Asterisks indicated statistical significance *, $P < 0.05$; **, $P < 0.01$, ***, $P < 0.001$, (Student's t-test). (H) Related to Figures 2C-E and S2E). Distribution of total nodule number per gram (g) root fresh weight of WT, *lsh-1*, *lsh2-1* and *lsh1-1/lsh2-1* plants grown in terragreen:sand at 28 days post inoculation with S. *meliloti*. A one-way Kruskal-Wallis rank sum test showed that total nodule number per plant is dependent on genotype (KW = 26.127, df = 3, $p < 8.97e-06$). Asterisk indicates a significantly different mean for *lsh1-1/lsh2-1* compared with WT, Dunn Test (95% confidence). (B-C and E-H) Box plots show median (thick line), second to third quartiles (box), minimum and maximum ranges (lines), and outliers (single points).

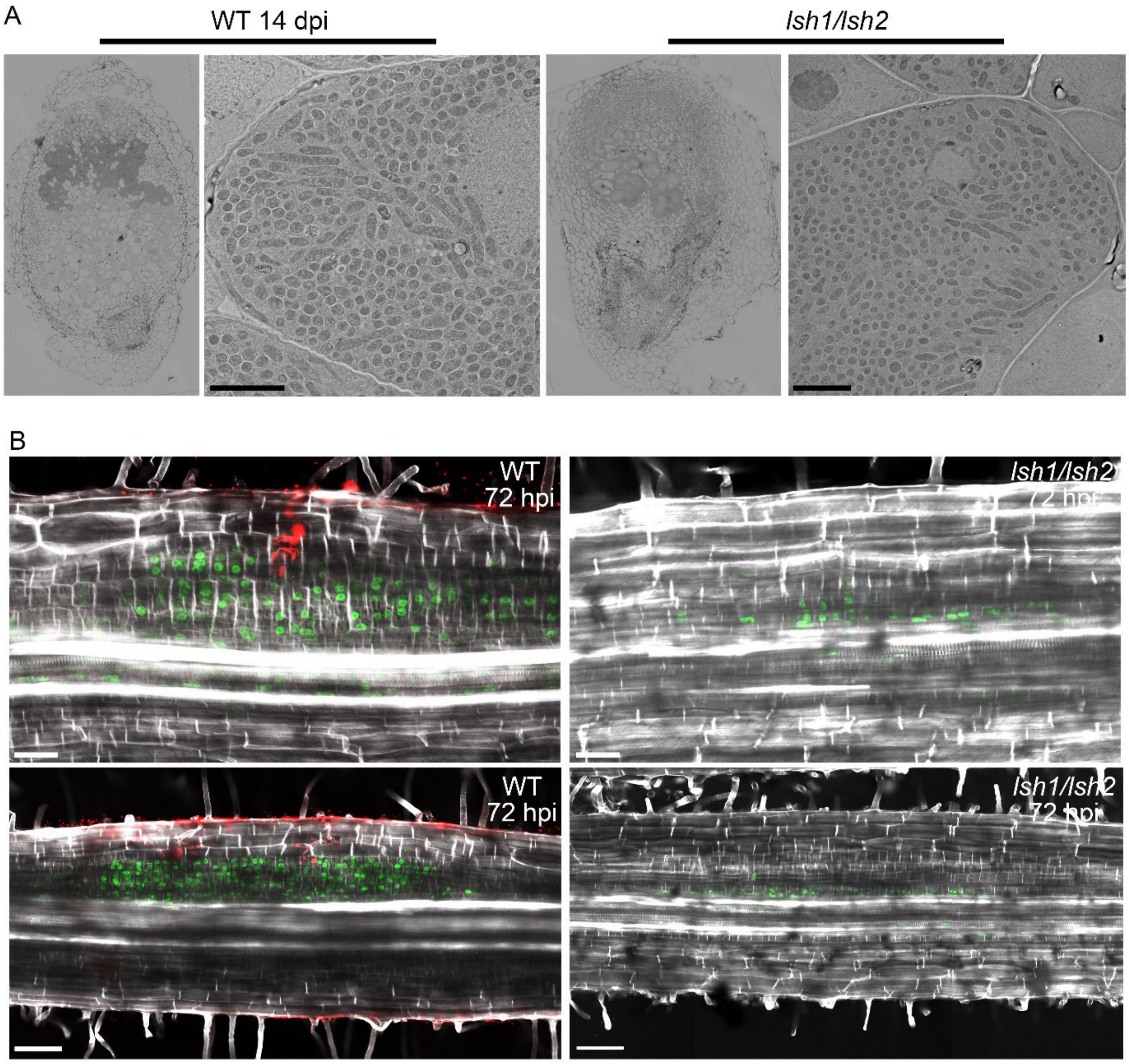


**Figure S3. *LSH* genes are required for the development of nodule primordia that can support bacterial colonization. Related to Figure 3.** (A) Representative light microscopy images of nodule section overviews (left panel) and transmission electron microscopy (TEM) images showing bacteroids (right panel) in WT and *lsh1/lsh2* at 14 dpi with *S. meliloti 2011*. Scale bars: 5 µm. (B) *LSH1* and *LSH2* are required for periclinal cell divisions during early nodule patterning. Related to Figure 3C. Optical sections of WT and *lsh1/lsh2* root sections 72 hours post spot-inoculation with *Sm2011-mCherry* bacteria. Top panel represents the overview images corresponding to the close ups in the bottom panel of Figure 3C. Bottom panel represents an additional set of WT and *lsh1/lsh2* samples. Infecting *Sm2011-mCherry* bacteria are labelled in red, fluorescent brightener staining (white) demarks cell walls and green EdU-labelled nuclei indicate DNA replication. Scale bars: 50 µm.

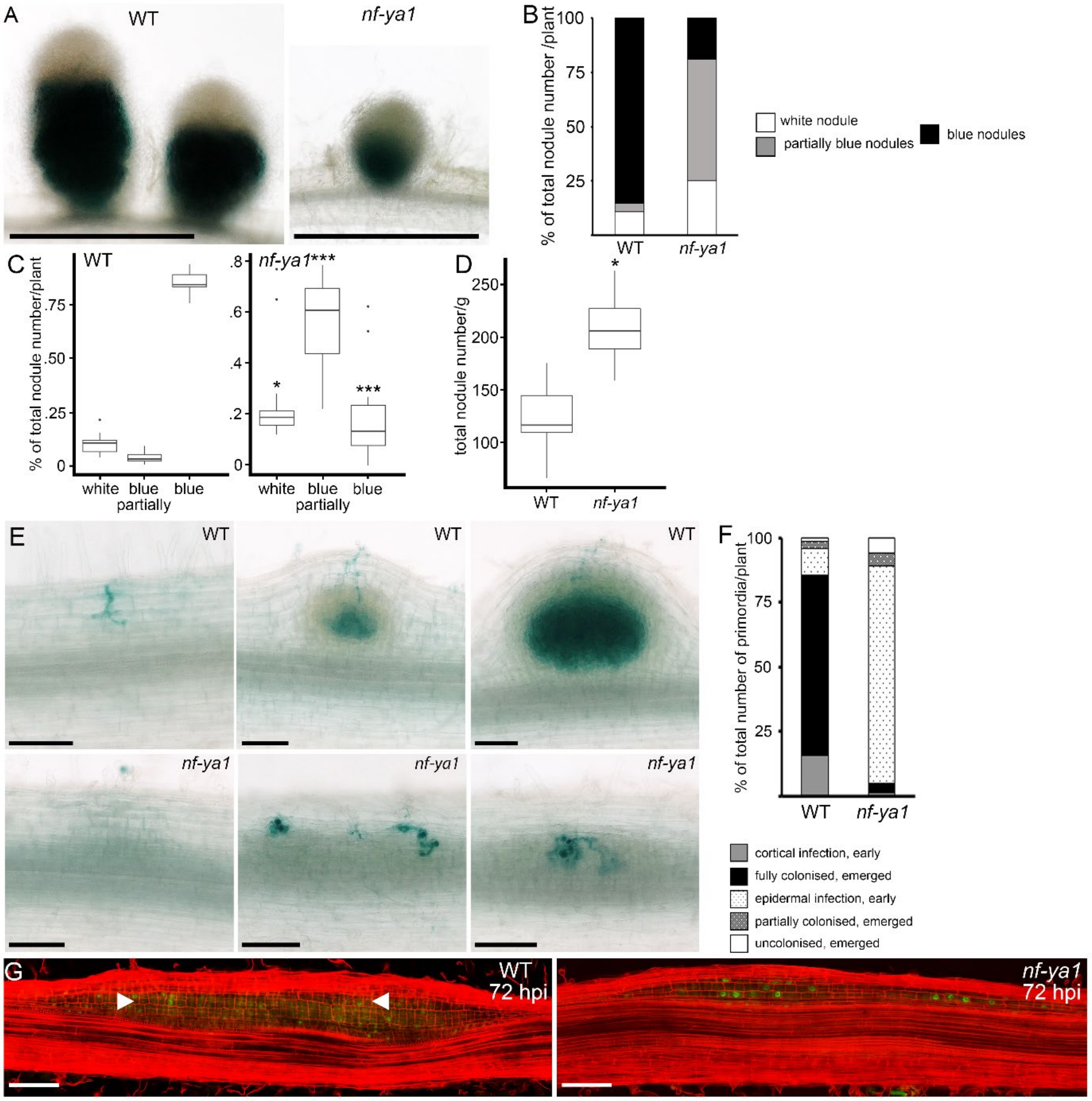


**Figure S4. *LSH1/LSH2* partly function through the cortical activation of *NF-YA1*. Related to Figure 5.** (A) Representative whole mount images of WT (jemalong) and *nfya1* in nodules 28 days post *S. meliloti* inoculation. GUS staining (blue) indicates the expression of the bacterial *pNifH* promoter as a marker for nitrogenase activity and nitrogen fixation. Scale bars: 500 µm. (B) Distribution of different nodule morphologies and N-fixation (*pNifH:GUS* staining) 28 days post *S. meliloti* inoculation depicted as percentage of the total number of nodules per plant and categorized as "white nodules" (white), *pNifH:GUS* expressing "partially blue nodules" (grey) and "blue nodules" (black) in WT (n = 12) and *nf-ya1* (n = 15) (for boxplots and statistical tests also see Figure S5C). (C) Distribution of nodule morphologies/types categorised as in (B) in WT (n = 12) and *nf-ya1* (n = 15) at 28 days post *S. meliloti* spray inoculation. Box plots show median (thick line), second to third quartiles (box), minimum and maximum ranges (lines), and outliers (single points). Student's t-tests showed that the distribution of the different nodule types is dependent on genotype; Asterisks indicated statistical significance *, $P < 0.05$; **, $P < 0.01$, ***, $P < 0.001$). (D) Related to Figures S5A-C. Distribution of total nodule number per gram (g) root fresh weight of WT and *nf-ya1* plants grown in terragreen:sand at 28 days post inoculation with S. *meliloti*. Asterisks indicated statistical significance *, $P < 0.05$; **, $P < 0.01$, ***, $P < 0.001$, (Student's t-test). (E) Representative images of WT and *nf-ya1* nodule primordia at different developmental stages, first initial divisions (left), multilayered (middle) and emerged primordia (right) observed 7 days post spray inoculation with rhizobial bacteria expressing *LacZ* (blue stain). Scale bars: 500 µm. (F) Related to Figure S5E. Distribution of bacterial colonization phenotypes observed in WT and *nf-ya1* plate-grown seedlings 7 dpi post *S. meliloti* inoculation, depicted as percentage of the total number of primordia per plant and categorized as "cortical infection in early multilayered primordia" (grey), "fully colonized emerged primordia" (black), "epidermal infection in early multilayered primordia" (white, hashed), "partially colonized emerged primordia" (grey hashed), and "uncolonized emerged primordia" (white) in WT ($n = 23$) and *nf-ya1* ($n = 28$). (G) Optical sections of WT ($n=14$) and *nf-ya1* ($n=29$) roots at 72 hours post spot inoculation with *S. meliloti*. Propidium iodide staining (red) demarks cell walls and EdU-labelled nuclei (green) indicate DNA replication. White arrowheads indicate periclinal cell divisions in the middle cortex. Scale bars:100 µm.

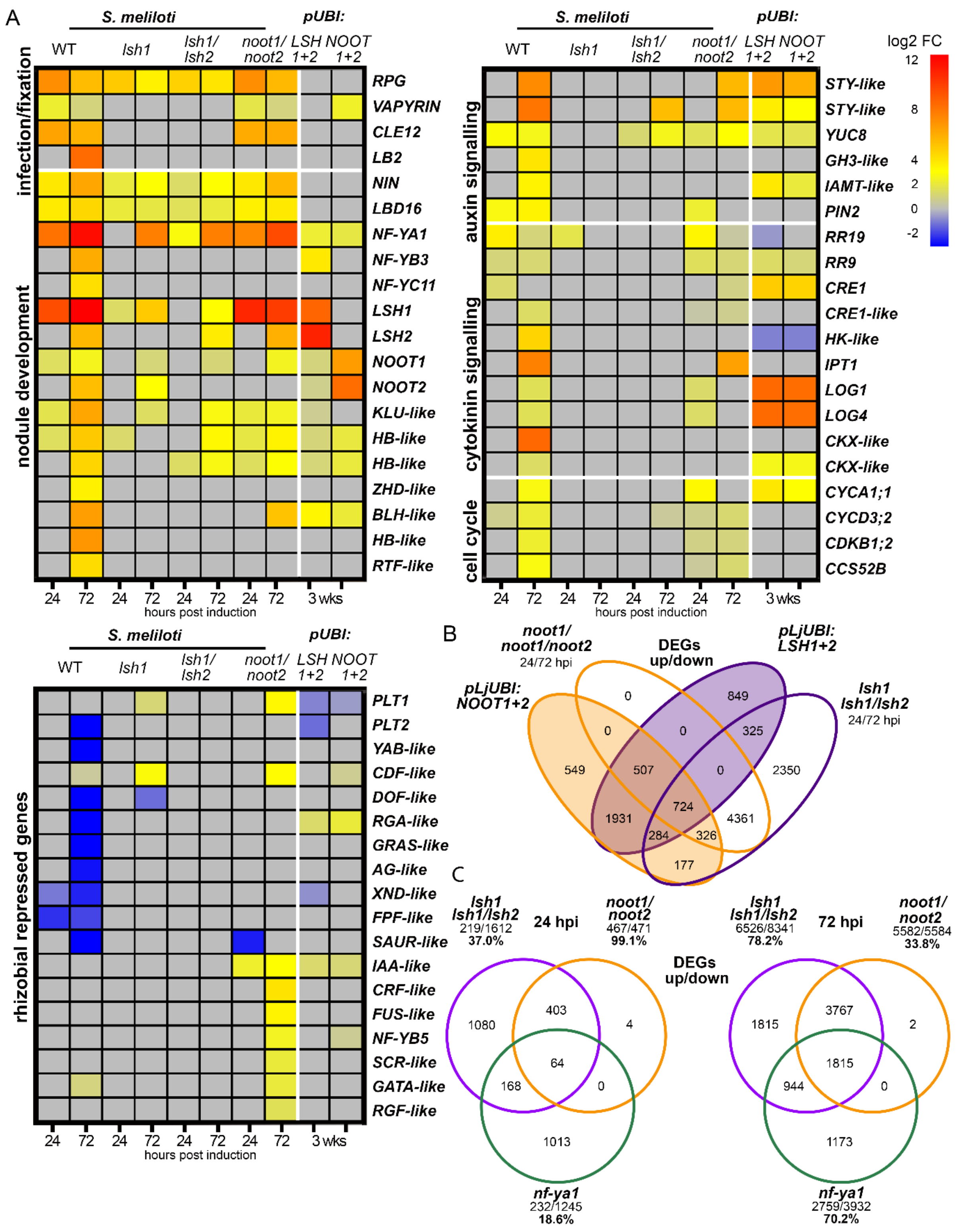


**Figure S5.** ***LSH1/LSH2*** **promote the expression of and act together with** ***NOOT1/NOOT2*** **in the same regulatory pathways. Related to Figure 6** (A) Related to Figures 6B-C also see Data S1 for gene identifiers and annotations. Heatmap of selected functional groups of genes with differential expression patterns in WT, *lsh1*, *lsh1/lsh2* and *noot1/noot2* root sections at 24 and 72 hours post spot inoculation with *S. meliloti* and in response to combined ectopic expression of *LSH1* and *LSH2* (*pUBI:LSH1+LSH2*) or *NOOT1* and *NOOT2* (*pLjUBI:NOOT1+NOOT2*) compared to empty vector control in 3-week old WT (jemalong) hairy roots under non-symbiotic conditions. Fold changes compared to controls are depicted in log2 scale with the significance threshold of p-value < 0.05. (B) Related to Figures 6B-C, also see Data S1. Comparison between all differentially expressed genes dependent on *lsh1* and/or *lsh1/lsh2* and *noot1/noot2* and genes differentially expressed in response to ectopic expression of *LSH1* and *LSH2* combined or *NOOT1* and *NOOT2* combined under the constitutive *LjUBI* promoter under non-symbiotic conditions. Genes with $\log_2$ fold changes of ≥ +/-1, p-value < 0.05 were included in this analysis. (C) Related to Figures 5B-C and 6B-C, also see Data S1. Comparison between all differentially expressed genes dependent on *lsh1* and/or *lsh1/lsh2* (purple circle), *noot1/noot2* (orange circle) and *nf-ya1* (green circle) up and down regulated at 24 or/and 72 hpi in response to spot inoculation with *S. meliloti*. Genes with $\log_2$ fold changes of ≥ +/-1, p-value < 0.05 were included in this analysis.

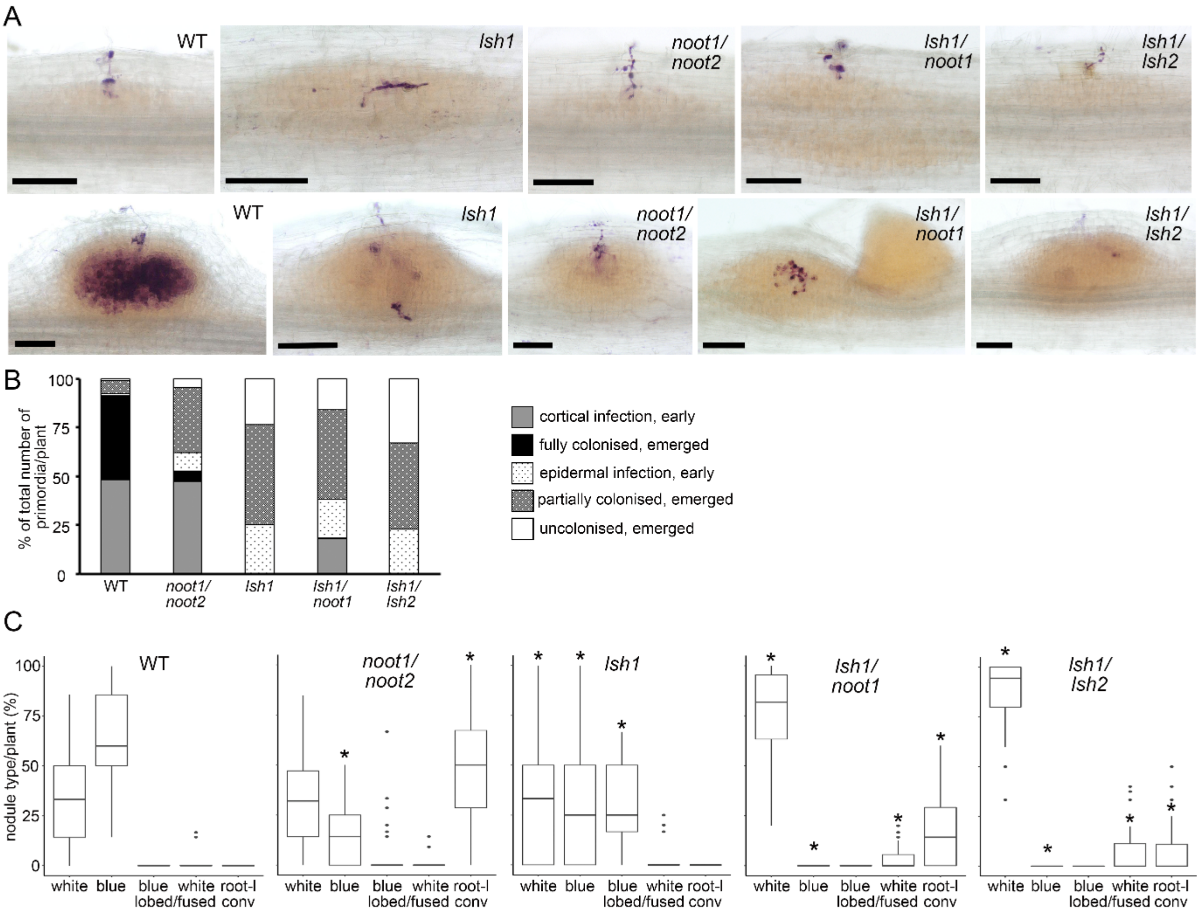


**Figure S6. Related to Figures 6 and 7. *LSH1/LSH2* and *NOOT1/NOOT2* function synergistically to confer nodule organ identity.** (A). Representative images of early-stage nodule primordia with first initial divisions (left), infection threads progressing through the outer cortical layers (middle) and emerged primordia (right) in WT, *lsh1*, *noot1/noot2*, *lsh1/noot1* and *lsh1/lsh2* observed 7 days post spray inoculation with *S. meliloti* expressing *LacZ* (magenta stain). Scale bars: 500 µm. (B) Distribution of bacterial colonization phenotypes observed in WT (n = 11), *lsh1* (n = 13), *noot1/noot2* (n = 12), *lsh1/noot1* (n = 15) and *lsh1/lsh2* (n = 12) at 7 dpi post inoculation with *S. meliloti* and depicted as percentage of the total number of primordia per plant categorized as "cortical infection in early multilayered primordia" (grey), "fully colonized emerged primordia" (black), "epidermal infection in early multilayered primordia" (white, hashed), "partially colonized emerged primordia" (dark grey hashed) and "uncolonized emerged primordia" (white). (C) Related to Figure 7A-B. Distribution of nodule morphologies/types categorised in "white", *pnifH:GUS* expressing "blue", "blue multilobed and/or fused", "white multilobed and/or fused" (indicated as lobed/fused) and "root-like conversions" (indicated as root-l conv) in percentage per plant for WT ( n = 37), *noot1/noot2* (n = 44), *lsh1* ( n = 33), ), *lsh1/noot1* (n = 35) and *lsh1/lsh2* (n = 40)

grown on plates at 21 days post spray inoculation with *S. meliloti*. Box plots show median (thick line), second to third quartiles (box), minimum and maximum ranges (lines), and outliers (single points). One-way Kruskal-Wallis rank sum tests showed that the distribution of nodule types is dependent on genotype (KW = 105.77 , df = 4, p = 2.2e-16 (white), KW = 129.21, df = 4, p = 2.2e-16 (blue), KW =105.83 , df = 4, p = 2.2e-16  (blue multilobed and/or fused), KW = 16.714, df = 4, p = 0.002196 (white multilobed and/or fused) and KW = 122.75, df = 4, p = 2.2e-16. Asterisks indicate significantly different means for *lsh1, noot1/noot2, lsh1/noot1* and *lsh1/lsh2* compared with WT, Dunn Test (95 % confidence).

| | n | keel fused | keel split | extra organs | P-value |
|---|---|---|---|---|---|
| WT | 20 | 20 | 0 | 0 | |
| % | | 100 | 0 | 0 | |
| *lsh1-2* | 22 | 1 | 14 | 7 | |
| % | | 4.55 | 63.64 | 31.82 | 4.09E-11 |
| *lsh2-1* | 20 | 20 | 0 | 0 | |
| % | | 100 | 0 | 0 | 1 |
| *lsh1-1/ lsh2-1* | 28 | 3 | 24 | 1 | |
| % | | 10.71 | 85.71 | 3.57 | 1.06E-10 |

**Table S1. Related to Figures 2A and S2D. Loss of *LSH1* function affects flower morphology.** Distribution of split and fused keel morphology and extra petals in flowers (as depicted in Figures 2A and S2D) of WT compared to *lsh1-2*, *lsh2-1* and *lsh1-1/lsh2-1* respectively, with P-values calculated using Fisher's exact test.

| construct expressed in *lsh1/lsh2* | n | number of nodules: white | partially blue | blue | total |
|---|---|---|---|---|---|
| empty vector control | 2 | 73 | 0 | 0 | 73 |
| % | | 100 | | | |
| *pLSH1:GFP-LSH1* | 10 | 118 | 16 | 21 | 155 |
| % | | 76.1 | 10.3 | 13.5 | |
| *pLjUBi:NF-YA1* | 6 | 170 | 27 | 12 | 209 |
| % | | 81.3 | 12.9 | 5.7 | |
| *pLSH1:NF-YA1 pLSH2:NF-YA1* | 10 | 175 | 54 | 8 | 237 |
| % | | 73.8 | 22.8 | 3.4 | |

**Table S2. Related to Figure 5E. Ectopic expression of *NF-YA1* partially rescues *lsh1/lsh2* nodule phenotype.** Distribution of "white", "partially blue" and "blue" nodules in absolute numbers and percentage of total nodule number per transformed hairy root system grown in terragreen:sand at 28 days post rhizobial spray inoculation.

| Name | Gene ID | Primer Sequence 5' to 3' | Primer Sequence 5' to 3' |
|---|---|---|---|
| **qRT-PCR** | | **Forward** | **Reverse** |
| *HH3* | *Medtr4g097170* | CCCTGGAACTGTTGCTCTTC | CCTGAGCAATTTCACGAACC |
| *LSH1* | *Medtr1g069825* | TCAAGAATCACCGTCCTCCTCTC | TGCCAAATTGGTCCAAGTACCG |
| *LSH2* | *Medtr7g097030* | AGAAACGAAAGCGTCCACCA | GAGTTGCACTTGCACCTTGT |
| **Genotyping** | | | |
| *Tnt1* | *Tnt1 transposon* | TCCTTGTTGGATTGGTAGCC | CAGTGAACGAGCAGAACCTGTG |
| *NF17203 lsh1-1* (WT) | *Medtr1g069825* | TGAAACCTGGAACATCTCTTG | CACCACTTTTTCTGCTACTTCA |
| *NF17203 lsh1-1* (rev mut) | *Medtr1g069825* | TGAAACCTGGAACATCTCTTG | Tnt1-F |
| *NF1304 lsh1-2* (WT) | *Medtr1g069825* | TGAAACCTGGAACATCTCTTG | CACCACTTTTTCTGCTACTTCA |
| *NF1304 lsh1-2* (fwd mut) | *Medtr1g069825* | TGAAACCTGGAACATCTCTTG | Tnt1-R |
| *NF14992 lsh2-1* (WT) | *Medtr7g097030* | GAATCATCGTCCACCTCTTTC | TACATTTGGAGAGGACAACACT |
| *NF14992 lsh2-1* (rev mut) | *Medtr7g097030* | GAATCATCGTCCACCTCTTTC | Tnt1-F |
| *NF2717 noot1-1* (WT) | Medtr7g090020 | GCAACAGAAACAACTAGCGA | GTGATGATGATACATGGAGTTG |
| *NF2717 noot1-1* (fwd mut) | Medtr7g090020 | Tnt1-F | TACACCAACCTTGAATCCAT |
| *NF5464 noot2-1* (WT) | *Medtr1g051025* | ACTCTAAGATCTCTCTCCCTTG | GCGGAAGAGAAGATTGGAAATA |
| *NF5464 noot2-1* (rev mut) | *Medtr1g051025* | ACTCTAAGATCTCTCTCCCTTG | Tnt1-F |

**Table S3. Primers used in this study. Related to STAR methods.**

| **L2 plasmids: GUS reporters** |
| --- |
| EC52371*_pL2B- R1-pAtUBI:KAN-R2-pMedtr1g069825(LSH1-4.51 kb):GUS-t-LSH1-3.13 kb-R3-pAtUBI:dsred-EC52371* |
| *EC52372_pL2B-R1-pAtUBI:KAN-R2-pMedtr7g097030(LSH2-4.98 kb):GUS-t-LSH2-2.21 kb-R3-pAtUBI:dsred-EC52372* |
| *EC11671_pL2B- R1-pAtUBI:KAN-R2-pMedtr7g090020(NOOT1-5.80 kb):GUS-t-NOOT1-2.71 kb-R3-pAtUBI:dsRed-EC11671* |
| *EC11672_pL2B-R1-pAtUBI:KAN-R2-pMedtr1g051025(NOOT2-5.48 kb):GUS-t-NOOT2-3.0 kb-R3-pAtUBI:dsRed-EC11672* |
| *EC43375_pL2B-R1-pMedtr1g056530(NF-YA1-5.99 kb):GUS-t-NF-YA1-1.63 kb-R2-pAtUBI:dsRed-EC43375* |
| **L2 plasmids: complementation and ectopic expression** |
| *EC43376_pL2B-R1-pMedtr1g069825(LSH1-4.51 kb)-GFP-LSH1-genomic-t-LSH1-3.13 kb-R2-AtUBI:dsred-EC43376* |
| *EC52316_pL2B-R1-pLjUBI:Medtr1g069825(LSH1-CDS)-t35S-R2-pLjUBI:Medtr7g097030(LSH2-CDS)-tOcs-R3-pAtUBI:dsred-EC52316* |
| *EC52380_pL2B-R1-pLjUBI:GFP-Medtr7g090020(NOOT1-CDS)-t35S-R2- pLjUBI:GFP-Medtr1g051025(NOOT2-CDS)-tOcs-R3-AtUBI:dsred-EC52380* |
| *EC52395_pL2B-R1-LjUBI:Medtr1g056530(NF-YA1-CDS)-tOcs-R2-AtUBI:dsred -EC52395* |
| *EC59515_pL2B-R1-pMedtr1g069825(LSH1-4.51kb):Medtr1g056530(NF-YA1-CDS)-t-LSH1-3.13 kb-R2-pMedtr7g097030(LSH2-4.98 kb):Medtr1g056530(NF-YA1-CDS)-t-LSH2-2.21kb-R3-pAtUBI:dsred-EC59515* |
| *EC11680_pL2B_R1-pAtUBI:KAN-R2-AtUBI:dsred-EC11680* |

**Table S4. Constructs used in this study. Related to STAR methods.**
